# Control of the *Azolla* symbiosis sexual reproduction: ferns to shed light on the origin of floral regulation?

**DOI:** 10.1101/2020.09.09.289736

**Authors:** Laura W. Dijkhuizen, Badraldin Ebrahim Sayed Tabatabaei, Paul Brouwer, Niels Rijken, Valerie A. Buijs, Erbil Güngör, Henriette Schluepmann

## Abstract

*Azolla* ferns and the filamentous cyanobacteria *Nostoc azollae* constitute a model symbiosis that enabled colonization of the water surface with traits highly desirable for development of more sustainable crops: their floating mats capture CO_2_ and fixate N_2_ at high rates phototrophically. Their mode of sexual reproduction is heterosporous. Regulation of the transition from vegetative to spore-forming phases in ferns is largely unknown, yet a pre-requisite for *Azolla* domestication, and of particular interest since ferns represent the sister lineage of seed plants.

Far-red light (FR) induced sporocarp formation in *A. filiculoides*. Sporocarps obtained, when crossed, verified species attribution of Netherlands strains but not Iran’s Anzali lagoon. FR-responsive transcripts included CMADS1 MIKC^C^-homologues and miRNA-controlled GAMYB transcription factors in the fern, transporters in *N*.*azollae*, and ycf2 in chloroplasts. Loci of conserved miRNA in the fern lineage included miR172, yet FR only induced miR529 and miR535, and reduced miR319 and miR159.

Suppression of sexual reproduction in both gametophyte and sporophyte-dominated plant lineages by red light is likely a convergent ecological strategy in open fields as the active control networks in the different lineages differ. MIKC^C^ transcription factor control of flowering and flower organ specification, however, likely originated from the diploid to haploid phase transition in the homosporous common ancestor of ferns and seed plants.

## INTRODUCTION

Ferns from the genus *Azolla* thrive floating on freshwater. They require no nitrogen whilst growing even at the highest rates due to an unusual symbiosis with the cyanobacterium *Nostoc azollae*, which fixates dinitrogen phototrophically (Brouwer *et al*., 2017). They are a model plant symbiosis allowing to study important traits: for example, highly efficient N_2_ fixation imparted by a microbial consortium, building of substrate mats with CO_2_ draw-down (Speelman *et al*., 2009), and adaptations to the aquatic floating lifestyle. Representative of the Monilophyte lineage moreover, their study is of interest as a sister lineage to seed plants to inform about the origin of seed plant traits. Previously used as biofertilizer, they now are considered a candidate crop for sustainable high-yield production of plant protein in subsiding delta regions but have never been domesticated (Brouwer *et al*., 2014a, 2018). What controls their sexual reproduction is completely unknown yet central to containment, propagation, breeding schemes or the establishment of biodiversity and germplasm banks.

*Azolla* ferns are heterosporous with diploid sporophytes generating two types of sori, mostly called sporocarps (Nagalingum *et al*., 2006; Coulter, 1910). Unlike homosporous ferns, the haploid phase of *Azolla* ferns, the gametophytes, are not free-living but contained inside the sporocarps. Unlike seed plants, gametophytes do not develop inside the sporocarps when these are still on the sporophytes. Therefore, fertilization is not guided by structures on the sporophyte such as flowers in angiosperms. *Azolla* sporocarps typically detach when mature and sink to the sediment of ditches; they constitute resting stages with nutrient reserves that survive drying or freezing and, thus, are key for dissemination practices (Brouwer *et al*., 2014).

The developmental transition to reproductive growth of the sporophyte (RD) occurs when shoot apices of the sporophytes will form initials for sporocarps in addition to branches, leaves and roots. RD is not well studied in ferns. It may be homologous to the transition in angiosperms of shoot apical meristems (SAM) to inflorescence meristems: the meristem is switching to making branches with leaves and sporocarps whereby the making of sporocarps is ancestral to the making of leaves (Vasco *et al*., 2016). Gene expression in lycophytes and ferns of Class III HD-Zip transcription factors (C3HDZ) was observed in both leaves and sporangia even though ferns and lycophyte fossil records have shown that the lineages evolved leaves independently. Vasco *et al*. (2016), therefore, suggested that the common C3HDZ role in sporangium development was co-opted for leaf development when leaf forms, the micro and megaphylls, evolved in lycophytes and ferns, respectively.

In the SAM transition to RD of the model angiosperm *Arabidopsis thaliana* (Arabidopsis), the LEAFY transcription factor (TF) complex induces floral homeotic genes (Sayou *et al*., 2016). LEAFY is conserved in plant lineages, but LEAFY does not promote formation of reproductive structures in seed-free plants including ferns where, nonetheless, the protein was reported to maintain apical stem cell activity (Plackett *et al*., 2018). The pattern emerging, therefore, is that the LEAFY complex is not among those that were already in place to link exogenous cues to the transition of SAM to RD in the common ancestor of seed plants and ferns.

In many angiosperms, MADS-box TF of MIKC^C^-type form complexes that integrate endogenous and exogenous cues turning SAM into RD (Theißen *et al*., 2018). In Arabidopsis, for example, gibberellic acid (GA)-dependent and photoperiod/temperature signaling is integrated by MIKC^C^ from the FLC and SVP clades that, as predicted by Vasco *et al*. (2016), are floral repressors. Consistent with separate transitions for inflorescence and flower meristems in Arabidopsis, the MIKC^C^ floral integrators act on another, SOC1, that in turn induces the expression of MIKC^C^ floral homeotic genes. Homeotic genes of potential relevance for ferns specify ovules, and stamens: these modules could be more ancient than those specifying carpels and sepals. Specification of ovules/stamens in angiosperms, however, is unlikely homologous to that of mega- and microsporocarps in *Azolla* ferns because heterospory evolved independently in ferns and angiosperms (Sessa & Der, 2016).

In addition to MIKC^C^ TF, micro RNA (miR) are known to control phase transitions in angiosperms, including RD. The miR156/172 age pathway controls flowering: miRNA156 decreases with the age of plants, and thus also its repression of the targets *SQUAMOSA PROMOTER BINDING-LIKES* (*SPL*). In the annual Arabidopsis, SPL promote expression of mir172 that targets another type of floral repressor, the AP2/TOE TF (Hyun *et al*., 2019). In the perennial *Arabis alpina*, the MIKC^C^ FLC homologue and miR156 regulate expression of *SPL15* thus integrating temperature and age-dependent RD. Moreover, AP2/TOE and NIN TF are known to mediate repression of *SOC1* in Arabidopsis on high nitrate (Gras *et al*., 2018; Olas *et al*., 2019). The MIKC^C^ SOC1 is further known to control levels of miR319 that inhibits TCP protein functions in the FT-FD complex mediating photoperiod control of RD (Lucero *et al*., 2017; Li et al., 2020). Inextricably, therefore, transition to RD is linked to conserved miRNA modules in angiosperms but we know little of miRNA loci in ferns (Berruezo *et al*., 2017; You *et al*., 2017).

*Azolla* ferns and Arabidopsis were shown to share responses to nitrogen starvation, suggesting that the ancestor of both had already evolved some of the responses to exogenous cues we know from angiosperms (Brouwer *et al*., 2017). GA applications to *Azolla* ferns, however, did not induce RD in *Azolla*, neither did stress treatment (Kar *et al*., 2002). Systematic research of the symbiosis transition to RD is lacking. Fern host and symbiont development are coordinated during the induction of sporocarps: mobile filaments of *N. azollae* generated in the shoot apical colony move into the developing sporocarp initials in the shoot apex (Figure 1). The host SAM developmental transition, therefore, may be influenced by activities of the *N. azollae* apical colony residing at the SAM (Figure 1A). Alternatively, *N. azollae* development may be controlled by secretions from specialized host trichomes in both the SAM, the developing leaf and sporocarp initials (Cohen *et al*., 2002; Perkins & Peters, 2006; Figure 1B and 1C). Transition to RD will be best characterized by profiling expression of the fern host and the symbiont simultaneously: we expect elaborate communication given that both genomes co-evolved and that *N. azollae* is an obligate symbiont with an eroded genome (Ran et al., 2010; Dijkhuizen et al., 2018; Li et al. 2018).

**Figure 1.**
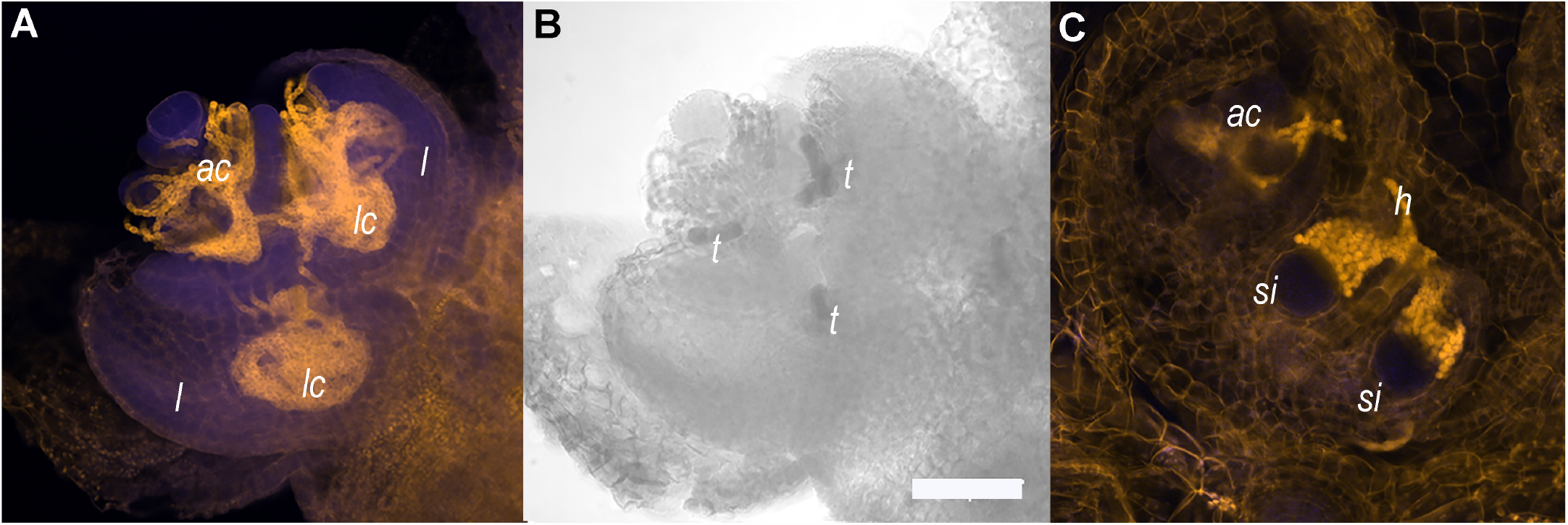
Shoot apices of *A. filiculoides* sporophytes contain the apical *N. azollae* colony, trichomes and sporocarp initials. **(A)** Shoot apex with apical *N. azollae* colony (*ac*), outer leaves were removed for better visualization of developing leaf lobes (*l*) and bacteria. **(B)** the same shoot apex as in (A) with trichomes (*t*) revealed by transmitting light absorption surrounded by the apical colony and in areas destined to form leaf cavities (*lc*); Bar: 50 μm. **(C)** *N. azollae* hormogonia (*h*) invading the sporocarp initials (*si*). Sporophytes were stained with propidium idiode (Truernit *et al*., 2008), then fluorescence detected using 405 nm activation beam, and emission <560 nm filter for the yellow channel and 505-530 nm for the blue channel with the Confocal Laser Scanning microscope Zeiss LSM5 Pascal.

Here we identify external cues that induce or inhibit transition to RD in *A. filiculoides* under laboratory settings. We test the viability of reproductive structures induced in different fern accessions by crosses, define species attributions and support the results obtained with phylogenetic analyses of the accessions. To reveal endogenous mechanisms mediating transition to RD, we profile transcript accumulation by dual RNA-sequencing comparing sporophytes grown with and without far-red light, at a time point when the reproductive structures are not yet visible. We assign induced MIKC^C^ and MYB TF to known clades using phylogenetic analyses. In addition, we profile small RNA from the symbiosis, then identify miRNA and their targets in the fern lineage. Finally, we discover regulatory modules of conserved miRNA/targets involved during induction of RD in ferns.

## RESULTS

### Far-red light induces, while nitrogen in the medium represses, the transition to sexual reproduction in *A. filiculoides*

A young fern growing at low density typically spreads over the water surface (Figure 2A). In contrast, a fern that underwent transition to reproductive development in dense canopies formed a three dimensional web characteristic of *Azolla* mats (Figure 2B). Unlike in the glasshouse, *A. filiculoides* ferns never sporulated under tube light (TL, Supplemental Figure 1A); TL did not emit any photons in the far-red range (FR, Supplemental Figure 1A). We, therefore, added far-red LED (FR, Supplemental Figure 1B) during the entire 16 h light period. This initially resulted in the elongation of the stem internodes of the fern (Figure 2C) and, after 2-3 weeks, in the formation of the sporocarps (Figure 2D). Sporocarps were positioned as pairs on the branch side of branch points and were predominantly arranged as a mega- and microsporocarp pair (Figure 2E).

**Figure 2.**
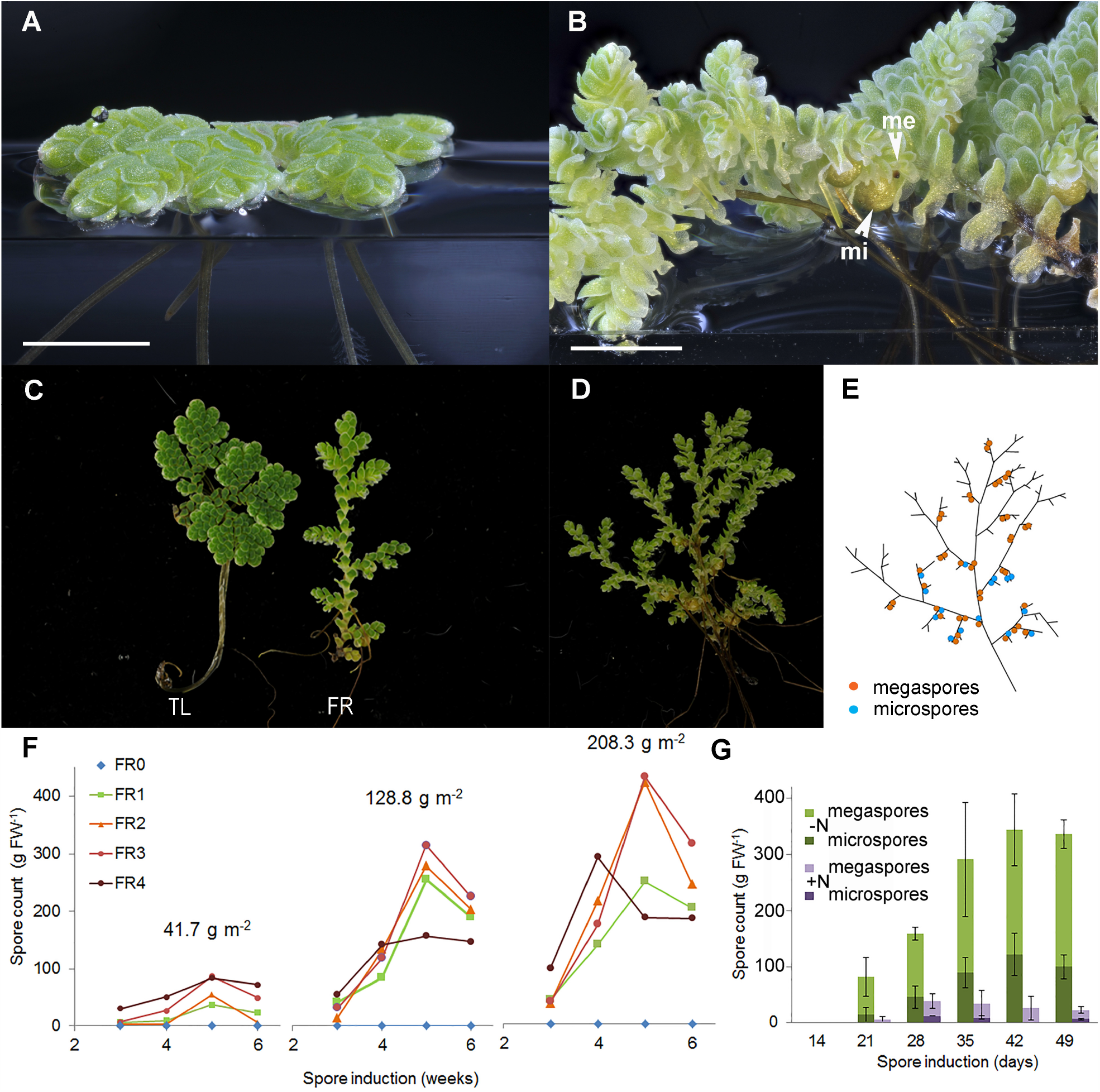
Far-red light and nitrogen affect the reproductive phase transition in *A. filiculoides*. **(A)** Vegetatively growing ferns. **(B)** Ferns in the reproductive phase with micro (mi) and megaspores (me). **(C)** Growth habit under TL and TL with far-red LED (FR). **(D)** A mature sporophyte and **(E)** its schematic representation: a root is formed at each branch point, and sporocarp pairs are found close-by on the branch side of the branch point but not at all branch points. **(F)** Sporulation frequencies at increasing plant density (41.7, 128.8 and 208.3 g m^-2^ dry weight) and with increasing red to far-red light ratios (FR0-FR4). The red to far-red ratios were 19.13 (FR0), 1.16 (FR1), 0.63 (FR2), 0.50 (FR3) and 0.31 (FR4); to obtain the FR4 ratio, the TL intensity was halved. Each data point represents the average of three independent continuous growth replicates. **(G)** Induction of sporulation without nitrogen (-N) or with 2 mM NH_4_NO_3_(+N) in the medium, data are averages from three replicates with standard deviations.

To test the optimum light for sporocarp induction, decreasing ratios of red to far-red LED were applied to ferns: sporophytes were first raised in TL light and then transferred to different red to far-red-light ratios whilst they were also maintained at three differing densities (Supplemental Table 2, Supplemental Figures 1C, 1D and 1E). Far-red light was required for sporocarp formation regardless of the density of the fern culture (Figure 2F). Increasing FR light whilst keeping a constant photosynthetic active radiation (PAR) yielded more sporocarps per fern mass. The sporocarps per fern mass increased with time of exposure to the FR up until five weeks and then decreased, presumably because ripe sporocarps detached. The sporocarps per fern mass, furthermore, increased with the density of the fern culture. Reducing the PAR at highest FR intensity (Figure 2F, FR4) revealed that PAR limits the production sporocarps at high FR light.

Adding 2 mM ammonium nitrate to the medium drastically reduced the spore count when inducing sporocarp formation with FR (Figure 2G). Both, ammonium and nitrate added individually inhibited sporocarp formation (data not shown), and inhibition explained why sporophytes without *N. azollae* kept on various nitrogen sources never sporulated.

The ratio of mega- to microsporocarp observed was approximately equal over nine weeks of induction with FR (Supplemental Figure 1F); and the marker *AzfiSOC1* (Brouwer *et al*., 2014) was maximally induced when sporocarps became visible four weeks after FR induction started (Supplemental Figure 1G).

### FR-induced sporocarps are viable which permits reciprocal crosses between strains to test phylogeny-derived species attributions

We next wondered whether the sporocarps induced with FR were viable. We randomly tested reciprocal crosses of the sequenced accession *A. filiculoides* Galgenwaard (Li *et al*., 2018), six further accessions from the Netherlands, one from Gran Canaria (Spain) and another from the Anzali lagoon (Iran) (Supplemental Table 1). All accessions could be induced to sporulate under FR, except Anzali. All sporulating accessions tested in crosses could be crossed suggesting that the ferns from the Netherlands and Spain were *A. filiculoides* (Table 1).

**Table 1.** Sporeling counts obtained from random crosses of *A. filiculoides* strains collected in The Netherlands and Spain (Gran Canaria).

| Megaspores: | Galgenwaard | Krommerijn | Hoogwoud | Gran<br>Canaria | Den<br>Bosch | Nijmegen | Nieuwebrug | Groningen |
| --- | --- | --- | --- | --- | --- | --- | --- | --- |
| Massulae : |  |  |  |  |  |  |  |  |
| Galgenwaard | >1000 | 17 |  | 1 |  | 13 |  | 2 |
| Krommerijn | 1 |  |  |  | 16 |  | 2 |  |
| Hoogwoud |  |  | 20 |  |  |  |  | 2 |
| Gran Canaria* | 17 |  |  |  | >40 | 1 |  |  |
| Den Bosch |  | 9 |  | 1 |  |  |  |  |
| Nijmegen |  | 12 |  | 33 |  |  | 2 |  |
| Nieuwebrug |  | 6 | 3 |  |  | 22 |  |  |
| Groningen |  |  | 1 |  |  |  | 5 |  |

\*crosses involving megaspores from the Gran Canaria strain reproducibly germinated late, generally after 5 weeks instead of 2-3 weeks.

To research the taxonomy of the accessions, intergenic regions of the chloroplast rRNA, used for phylogenetic studies previously (Madeira *et al*., 2016), were amplified and sequenced to compute phylogenetic relations. The regions trnL-trnF (Figure 3A), and trnG-trnR (Figure 3B) yielded similar trees confirming the close relationship of the *A. filiculoides* accessions from the Netherlands and Spain. The trees further revealed that the Anzali accession clustered along with two accessions from South America that do not cluster within the *A. caroliniana* group (Southern Brazil and Uruguay, Supplemental Table 1). Use of the nuclear ITS region for phylogeny reconstruction proved futile when amplifying the region on the *A. filiculoides* genome *in silico*: the *A. filiculoides* genome assembly contained over 500 copies of the ITS sequence, and the copies of ITS were variable within the same genome (Figure 3C). Variations within anyone genome were furthermore large when comparing variations between genomes of *Azolla* species (Supplemental Figure 3A).

**Figure 3.**
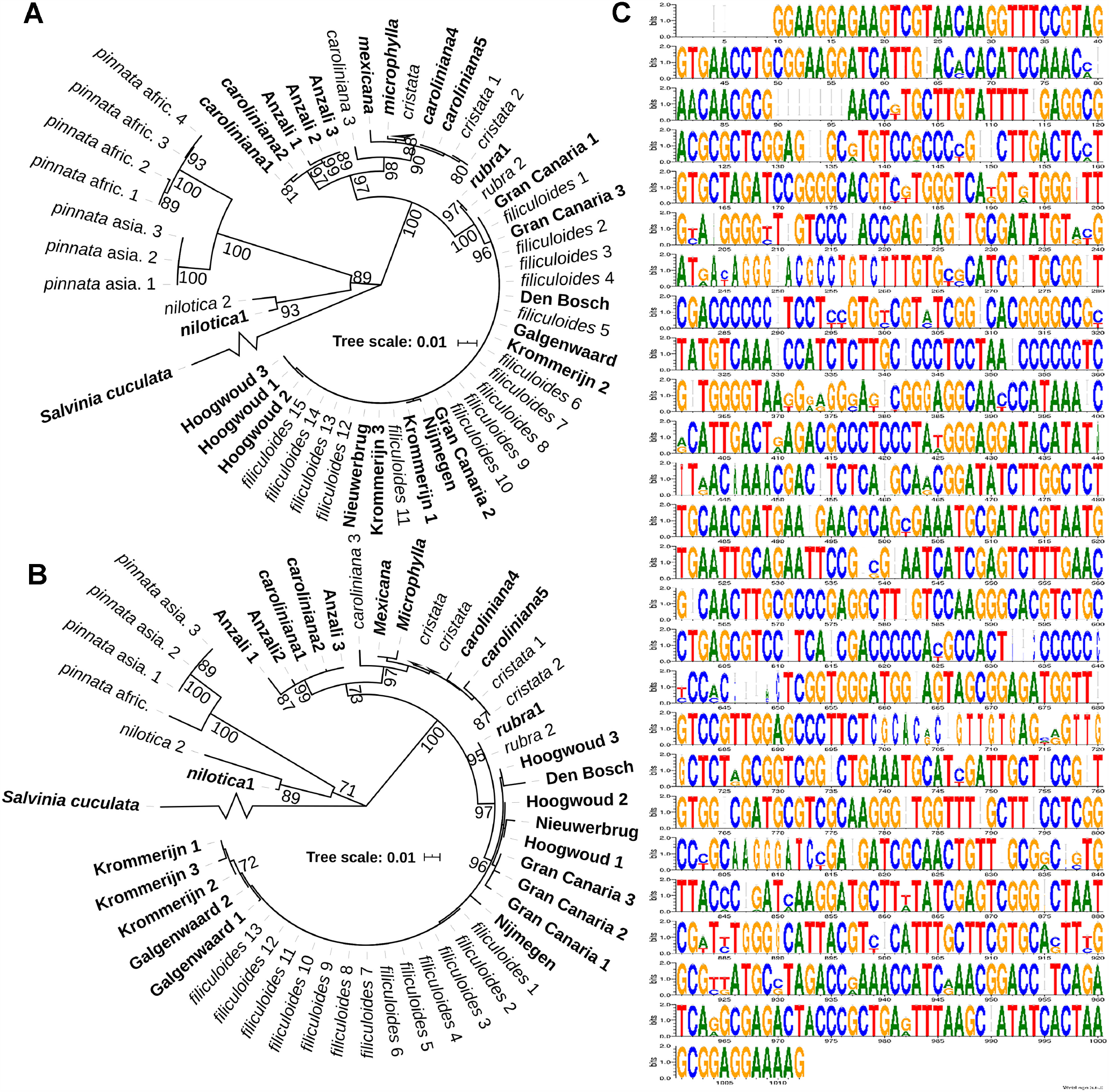
Chloroplast marker regions in strains from the Netherlands, Spain and Iran, and the rRNA intergenic regions of the *A. filiculoides* genome. **(A)** Phylogenetic tree of chloroplast regions trnL-trnF. The maximum likelihood trees were bootstrapped 1000x. All bootstrap values >0.70 are displayed. In bold: sequences from accessions in this study labelled after their collection site and sequences from the differing species extracted from genome shot-gun sequencing (Li *et al*., 2018). In regular type: sequences from the Madeira et al. (2016) study. **(B)** Phylogenetic tree of chloroplast regions trnG-trnR. **(C)** Sequence logo of all ITS1 gene regions in the *A. filiculoides* genome (Li *et al*., 2018). The gene regions were extracted and aligned (MEGA 7) then used for *in-silico* PCR. Results were visualized using WebLogo (vs 2.8.2 Crooks *et al*., 2004): at each position of the ITS, the height of the stack indicates the sequence conservation, while the height of symbols within the stack indicates the relative frequency of each base.

The dorsal side of the upper leaf lobes of the Anzali accession had two-celled, distinctly pointed, papillae unlike *A. filiculoides* that mostly, but not exclusively, have single-celled and rounded papillae (Supplemental Figure 3B). Consistent with its differing response to far-red light, and its phylogenetic position based on the chloroplast sequences, the Anzali accession, we conclude, is not *A. filiculoides*.

### Dual RNA sequencing by rRNA depletion profiles RNA from organelle, prokaryotic symbiont and fern nucleus

We nest wondered which molecular processes are altered during the symbiosis transition to RD. Ferns maintained on TL were transferred to TL with and without FR for a week in triplicate cultures, then collected 2 h into the 16 h light period snap frozen. Total RNA was then extracted, its plant rRNA depleted, and stranded sequencing libraries generated which were depleted for rRNA from gram-negative bacteria. Libraries were sequenced to obtain 12 to 60 million paired-end (PE) 50 b reads, quality filtered and trimmed. To align the reads obtained, nuclear and chloroplast genomes with annotations for the *A*.*filiculoides* Galgenwaard accession were available (Li *et al*., 2018) but not for its cyanobacterial symbiont.

The *N. azollae* Galgenwaard genome was assembled both from plants kept in the lab (*A. filiculoides* lab; Li et al., 2018) and plants taken from the original sampling location (*A. filiculoides* wild; Dijkhuizen et al. 2018). Both assemblies were from short-reads of the entire symbiosis metagenome and were fragmented but highly complete: over 95% based on single copy marker genes. Contigs aligning to the *N. azollae* 0708 reference (Ran *et al*., 2010) were well over 99.5% identical to each other (Figure 4A). This degree of nucleotide identity was well above the maximum mismatch of STAR-aligner’s default setting; PE reads were thus aligned to the Galgenwaard accession nuclear and chloroplast genomes, and the *N. azollae* 0708 genome, and read counts derived from their GFF annotation files. Alignments revealed a high proportion of uniquely mapped reads to the fern nuclear and *N. azollae* genomes suggesting efficient removal of rRNA (Figure 4B). The larger proportion of PE reads mapping to multiple loci in the chloroplast indicated that fern chloroplast rRNA was not as efficiently removed (Figure 4B), yet read counts per gene were still high because few genes are encoded in the chloroplast genome (84 without rRNA and tRNA genes). Read counts in genome features further revealed that sense alignments dominated and thus that the libraries were stranded (Figure 4C).

**Figure 4.**
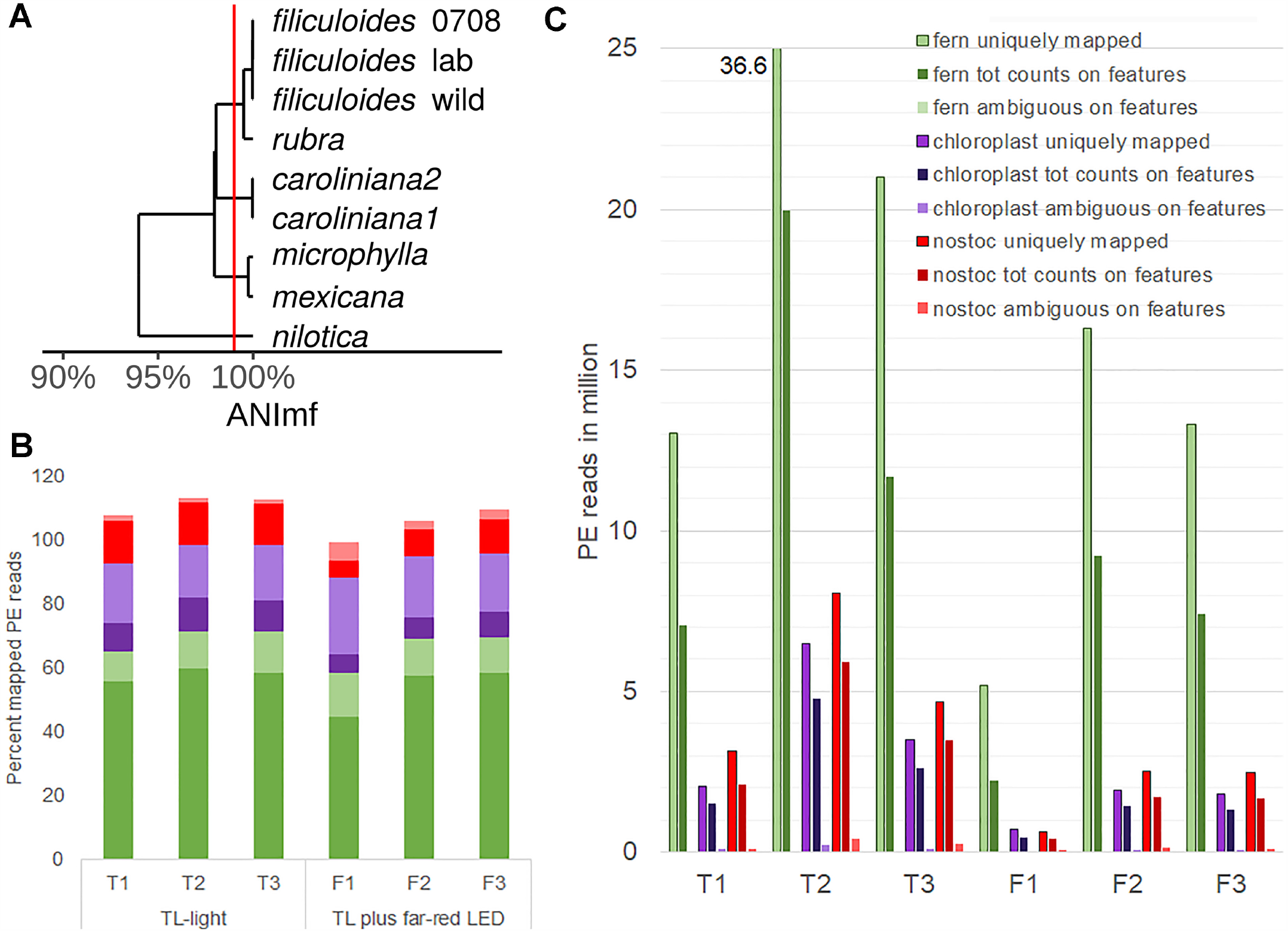
Profiling eukaryotic and prokaryotic RNA simultaneously in *A. filiculoides* (dual RNAseq). **(A)** Similarity of the *N. azollae* 0708 genome and Metagenome Assembled Genomes (MAGs) from *N. azollae* strains in *Azolla* species. Average Nucleotide Identity (ANI) was calculated with the dRep implementation of nucmer using the ANImf preset (Kurtz *et al*., 2004; Olm *et al*., 2017). Pairwise average ANI is shown in an UPGMA dendrogram (de Vries & Ripley, 2016; Wickham, 2011). Aligned MAGS covered over 90% of the reference *N. azollae* 0708. **(B)** Proportion (percent) of uniquely aligned PE reads (dark color) and PE reads aligning at multiple loci (light color) on the genomes of fern (green), fern chloroplast (purple) and *N. azollae 0708* (red). PE reads were from samples of sporophytes grown for a week on TL-light (replicates T1, T2, T3) or TL light with far-red LED (replicates F1, F2, F3). STAR-aligner was used in PE mode. **(C)** Yields of PE uniquely mapped compared with PE read counts on features and PE reads that STAR aligned ambiguously to several features. Features were specified by current general features files (GFF) of each of the fern (green), chloroplast (purple) and *N*.*azollae* 0708 (red) genomes.

Dispersion of sense read count per gene was low which permitted identification of differentially accumulating transcripts (defined as DEseq2 Padj <0.1; Love *et al*., 2014) comparing ferns on TL and TL with FR: 318 from the fern nucleus, 67 from *N. azollae* and five from the fern chloroplast (Supplemental Table 4). We next wondered whether the changes in transcript abundance reflect adaptations to light quality and the transition to reproductive development.

### FR triggers small transcriptional changes reflecting light-harvesting adaptations in chloroplasts and *N. azollae*, yet large changes in transporters and transposases of *N. azollae*

After a week exposure FR, increased chloroplast transcripts encoded ycf2 (log2 of 2.9), energizing transport of proteins into the chloroplast, and RNA polymerase subunit rpoA (log 2 of 1.3) (Supplemental Table 4, *A. filiculoides* Chloroplast). Decreased chloroplast transcripts included psbD, previously reported as responsive to light quality in plant lineages (Shimmura *et al*., 2017).

Consistent with the high levels of N_2_ fixation measured in *Azolla* ferns, the highest *N. azollae* read counts were in transcripts of the *nif* operon, genes encoding photosystem I and II proteins and ATP synthase subunits. In addition, read counts for *ntcA* and metabolic enzymes supporting high N_2_ fixation were high (Supplemental Table 5). Read counts from the *N. azollae* transcripts reflected the activity of the symbiont and we thus further analyzed the data for differential expression.

In *N. azollae*, far-red light led to accumulation of the PsbA transcript, but reduction of photosystem I reaction center subunit XII, geranylgeranyl hydrogenase ChlP, and phycobilisome rod-core linker polypeptide CpcG2 transcripts (Supplemental Table 4, *N. azollae*). These changes in light harvesting-related genes were small, however, compared to changes in hypothetical proteins of unknown function, transposases and highly expressed transporters (TrKA, MFS and ABC transporters). We conclude that a one-week induction of reproductive structures with far-red LED triggered few transcriptional changes reflecting light-harvesting adaptations in *N. azollae*. The larger differential accumulation of transporter transcripts reflects changes in metabolite trafficking and communication with the host fern.

### FR alters transcripts from MIKC^C^ and R2R3MYB TF related to those of the RD of seed plants

As in the case of *N. azollae* symbiont, the ferns largest changes in transcript accumulation under FR were in genes with unknown function, then in secondary- and lipid-metabolism (Supplemental Table 4, *A. filiculoides* nucleus). We noted the large induction of the SWEET-like transporter (Azfi_s0221.g058704 log2 7.5 fold), of transcripts related to cysteine-rich peptide signaling (Azfi_s0046.g030195 log2 7.1 fold; Azfi_s0745.g084813 log2 4.0 fold) and of transcripts from jasmonate metabolism. To examine the link between RD in seed plants and ferns, however, TF were analyzed in more depth. Transcripts of MIKC^C^ TF and of R2R3MYB TF were strikingly induced in sporophytes under FR (Table 2).

**Table 2.** Transcription factors with largest changes in transcript abundance when comparing sporophytes in TL with and without far-red LED.

| *A. filiculoides* locus | baseMean | Log2Fold Change | DESeq2 padj | Mercator 4.0 annotation | Closest Arabidopsis Homolog |
| --- | --- | --- | --- | --- | --- |
| **Azfi_s0028.g024032** | **9** | **7.22** | **0.001** | **15.5.14 MADS/AGL** | **AGL20/SOC1, MIKC<sup>c</sup>** |
| Azfi_s0113.g045874 | 155 | 6.41 | 0.002 | 15.5.2 R2R3MYB | AtMYB 32 (AT4G34990.1), R2R3MYB VIII-E* |
| Azfi_s0083.g038807 | 59 | 5.75 | 0 | 15.5.32 Basic Helix-Loop-Helix | EDA33, IND1, INDEHISCENT (AT4G00120.1) |
| Azfi_s0003.g007560 | 8 | 5.27 | 0.011 | 15.5.32 Basic Helix-Loop-Helix | FIT1 (regulates iron transport) |
| Azfi_s0003.g007559 | 34 | 3.49 | 0 | 15.5.32 Basic Helix-Loop-Helix | FIT1 (regulates iron transport) |
| Azfi_s0096.g043715 | 31 | 3.13 | 0.001 | 15.5.7 DREB subfamily A-2 of ERF/AP2 | AT5G18450.1 |
| **Azfi_s0003.g007710** | **1778** | **2.82** | **0** | **15.5.14 MADS/AGL** | **AGL6, MIKC<sup>c</sup>** |
| Azfi_s0015.g013719 | 111 | 2.57 | 0 | 15.5.2 G2-like, GARP | HHO2, NIGT1.2 |
| Azfi_s0112.g045798 | 66 | 2.2 | 0 | 15.5.51.1 NF-Y component NF-YA | AT5G12840.4 |
| Azfi_s0014.g013539 | 105 | 2.07 | 0.005 | **15.5.2 R2R3MYB** | **LOF2, R2R3MYB V\*** |
| Azfi_s0015.g014012 | 165 | 1.99 | 0.026 | 15.5.17 NAC domain | NAC025 |
| **Azfi_s0004.g008455** | **103** | **1.93** | **0.003** | **15.5.2 R2R3MYB** | **miRNA319 controlled GAMYB 33, R2R3MYB VII\*** |
| Azfi_s0132.g049213 | 63 | -2.35 | 0.004 | 15.5.32 Basic Helix-Loop-Helix | No hits |
| Azfi_s0182.g056462 | 10 | -4.46 | 0.049 | 15.5.3 HD-ZIP I/II | HB16 |

\*R2R3 MYB classification according to Jiang & Rao, 2020.

Prediction of the most induced TF transcript annotated as SOC1-like MIKC^C^ (Azfi_s0028.g024032, log2 7.2 fold) missed the K-domain because this TF was not expressed in vegetative sporophytes. Manual annotation allowed computation of a phylogenetic tree (Supplemental Figure 4): both induced *Azolla* MIKC^C^ were in the well-supported clade (95.5% bootstrap confidence) of fern sequences radiating separately from the angiosperm and gymnosperm MIKC^C^ clade that gave rise to AtSOC1, FLC, and the A,C,D and E-function floral homeotic genes (Supplemental Figure 4). Induction of the fern SOC1-paralogue was confirmed by RT-PCR (Supplemental Figure 2G). In Arabidopsis, AtSOC1 is known to control the steady state of miR319 that inhibits TCP TF functions necessary for the transition to an inflorescence meristem at the shoot apex (Lucero *et al*., 2017); the link may also operate in *Azolla* ferns since a TCP TF (Azfi_s0168.g054602) was induced in ferns on TL with far-red LED (Supplemental Table 4, *A. filiculoides* nucleus).

Transcripts of R2R3MYB TF could be assigned to families using data from Jiang & Rao, 2020 (Table 2). The highly induced class VIII-E TF (log2 6.1 fold) likely controls changes in secondary metabolism together with the similarly changed bHLH TF (Güngör *et al*., 2020), and therefore may not be linked to RD directly. In contrast, induced class V (log 2 2.1 fold) and VII (log 2 1.9 fold) TF are known to affect seed plant RD, with the GA pathway associated class VII GAMYB in seed plants known to be regulated by miR319.

Prominence of transcription factors in Table 2 suspected to controlling RD as in seed plants begged the question as to whether any RD pathways with conserved miRNA were conserved in the common ancestor of seed plants and ferns. This required to first characterize miRNA loci in the *A*.*filiculoides* symbiosis.

### Small RNA profiles are characteristic for fern nucleus, chloroplast or *N. azollae*

Short RNA reads were obtained from sporophytes after a week growth under TL without and with FR from the same cultures as those used for dual RNA sequencing. In addition, reads were obtained from sporophytes grown under TL with far-red LED on medium with 2 mM NH_4_NO_3_, and from sporophytes without *N. azollae* on medium with 2 mM NH_4_NO_3_. All samples were collected two hours into the 16 h light period. The sRNA sequencing library preparation chemistry did not distinguish pro-from eukaryote sRNA and therefore was inherently a dual profiling method. Reads obtained were aligned to concatenated genomes which yielded small RNA profiles characteristic for the fern nucleus, chloroplast and *N. azollae* (Figure 5; Supplemental Figure 5).

**Figure 5.**
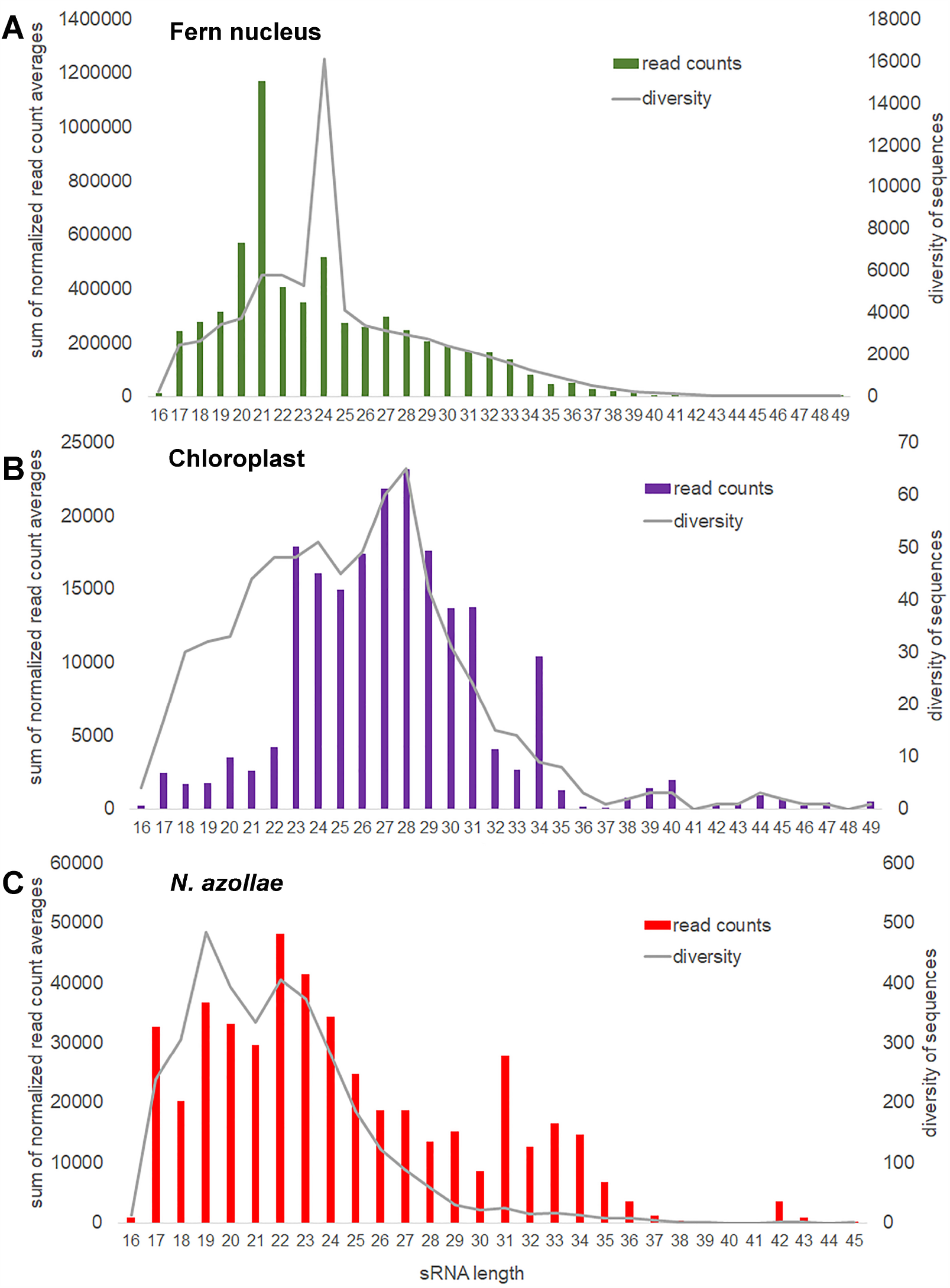
sRNA size distribution and diversity for each of the genomes in the *A. filiculoides* symbiosis. sRNA reads were mapped with 0 mismatches, no overhang, to the concatenated genomes; bam2fasta was then used to extract sRNA sequences aligning per genome, the fasta files obtained were then collapsed to compute read counts for each sRNA, the lists obtained were then joined to a read count matrix imposing at least 1 read per sRNA in each of the three replicate sample types: f (far-red), t (no far-red) and fn (far-red and nitrogen in the medium). sRNA counts were then normalized to the total read counts per sample for that genome. **(A)** Fern nucleus. **(B)** Chloroplast, included samples from sporophytes without cyanobacteria in the analysis. **(C)** *N. azollae*, read count files joined imposing at least 10 reads per sRNA in every sample.

The most abundant sRNA encoded in the fern nucleus were 21, 20 and 24 nt long, but the most diverse were 24 nt long, consistent with abundant 21 nt miRNA discretely cut from a hair-pin precursor by dicer, and the 24 nt siRNA generated by random processes (Figure 5A). The chloroplast most abundant sRNA of 28 nt was also the most diverse, but the chloroplast sRNA have an unusual peak of sRNA of low diversity at 34 nt (Figure 5B). The sRNA encoded in *N. azollae* are generally shorter than those of the chloroplast, yet, they have a conspicuous peak of low diversity at 42 nt length, consistent in length with CRISPR RNA (Figure 5C).

### Conserved miRNA in *A. filiculoides* include miR172 and 396, but only miR319 and 159 were decreased in response to FR, whilst miR529 and 535 were increased

Fern genome-encoded sRNA reads of 20-25 nt length were used for miRNA predictions allowing a hairpin of maximally 500 bp. Loci predicted were then verified against miRNA criteria (Axtell & Meyers, 2018) as, for example, in the case of the miRNA 172 locus Azfi_s0021g15800 (Figure 6): the locus was expressed as a pre-miRNA (Figure 6A, RNAseq all conditions) and both the miRNA and miRNA* are found in replicate samples and cut precisely with two base overhangs (Figure 6B and 6C, sRNAseq). The miR172 reads were derived from the fern since they were also found in sporophytes without *N. azollae* (Figure 6, RNAseq -*N. azollae*). The locus encoded a stable long hairpin which when folded released -65.76 kcal mol^-1^ at 37°C. MiR172 targets in seed plants include the AP2/TOE transcription factors, and alignment of the target sites to all annotated AP2/TOE transcripts from *A. filiculoides* revealed target sites for the AzfimiR172 (Supplemental Figure 6). The miR156a locus was similarly verified (Supplemental Figure 7).

**Figure 6.**
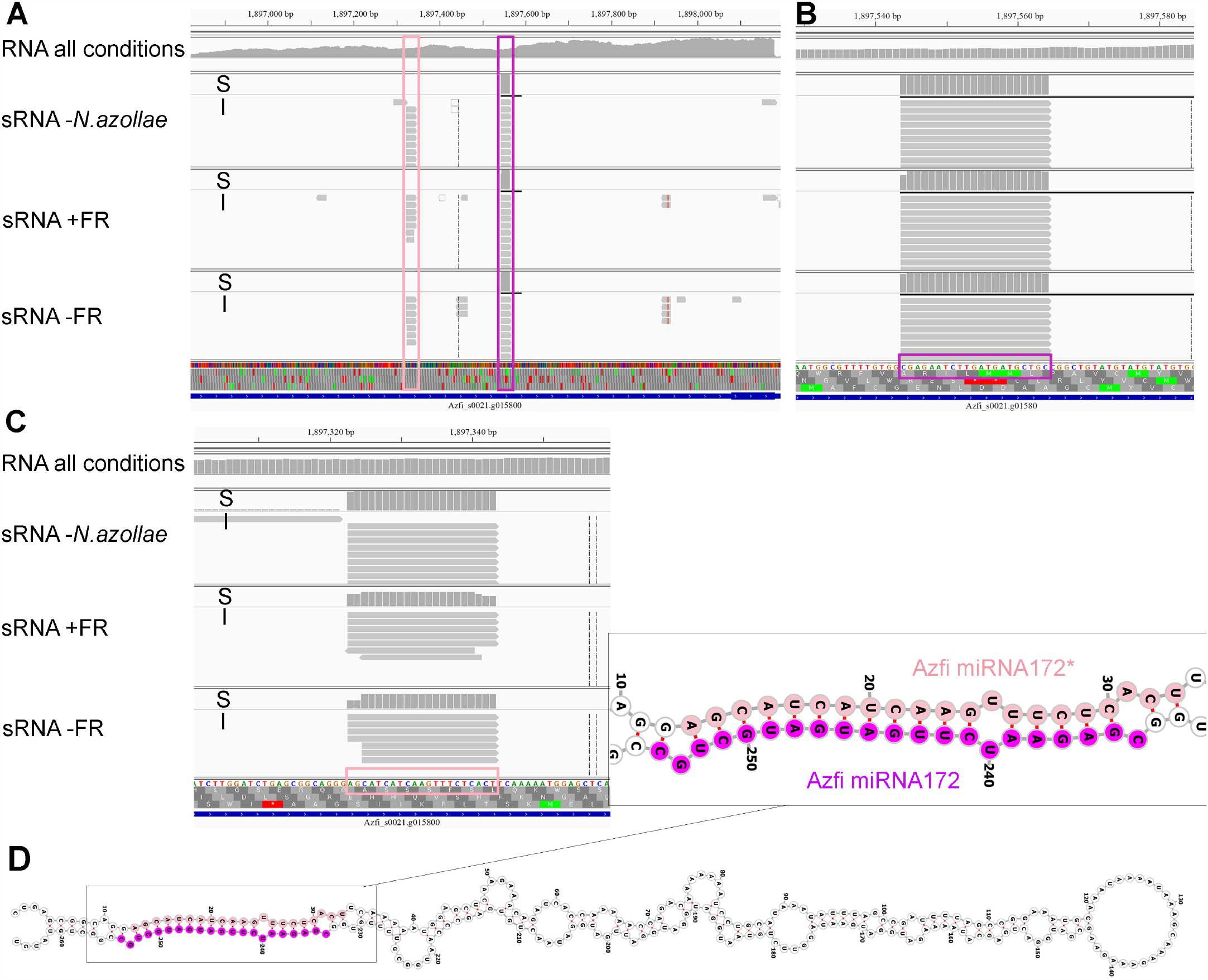
The miRNA 172α locus in *A*.*filiculoides*. Alignments at the Azfi_miRNA172α locus are visualized using the Integrative Genomics Viewer (Thorvaldsdóttir *et al*., 2013); S, read summary; i, individual reads. **(A)** Reads from pre-miRNA at the miR172α locus. RNAseq all conditions, all reads pooled from the diel RNAseq experiment in (Brouwer *et al*., 2017). **(B)** Reads of the miR172 in sRNA libraries including those of ferns without *N*.*azollae* (sRNAseq-*N*.*azollae*), ferns grown with and without far-red LED (sRNAseq+FR, sRNAseq TL-FR). **(C)** Reads of the miR172*. **(D)** Azfi_miRNA172α hairpin folded using Vienna RNAfold (Gruber *et al*., 2008; predicts -104,07 kcal mol^-1^ at 22 ^°^ C upon folding); miRNA (purple) and miRNA* (pink) stem arms are shown with 3’overhangs.

Mature miRNA were compared in miRbase (Kozomara *et al*., 2019) which identified 11 of the conserved miRNA families of 21 nt in the *A. filiculoides* genome and thus in the fern lineage (Figure 7A). The list included miR396, as well as miR172, not yet confirmed in other seed-free plant lineages. MiRNA predictions further uncovered a number of novel miRNA (miRNF).

**Figure 7.**
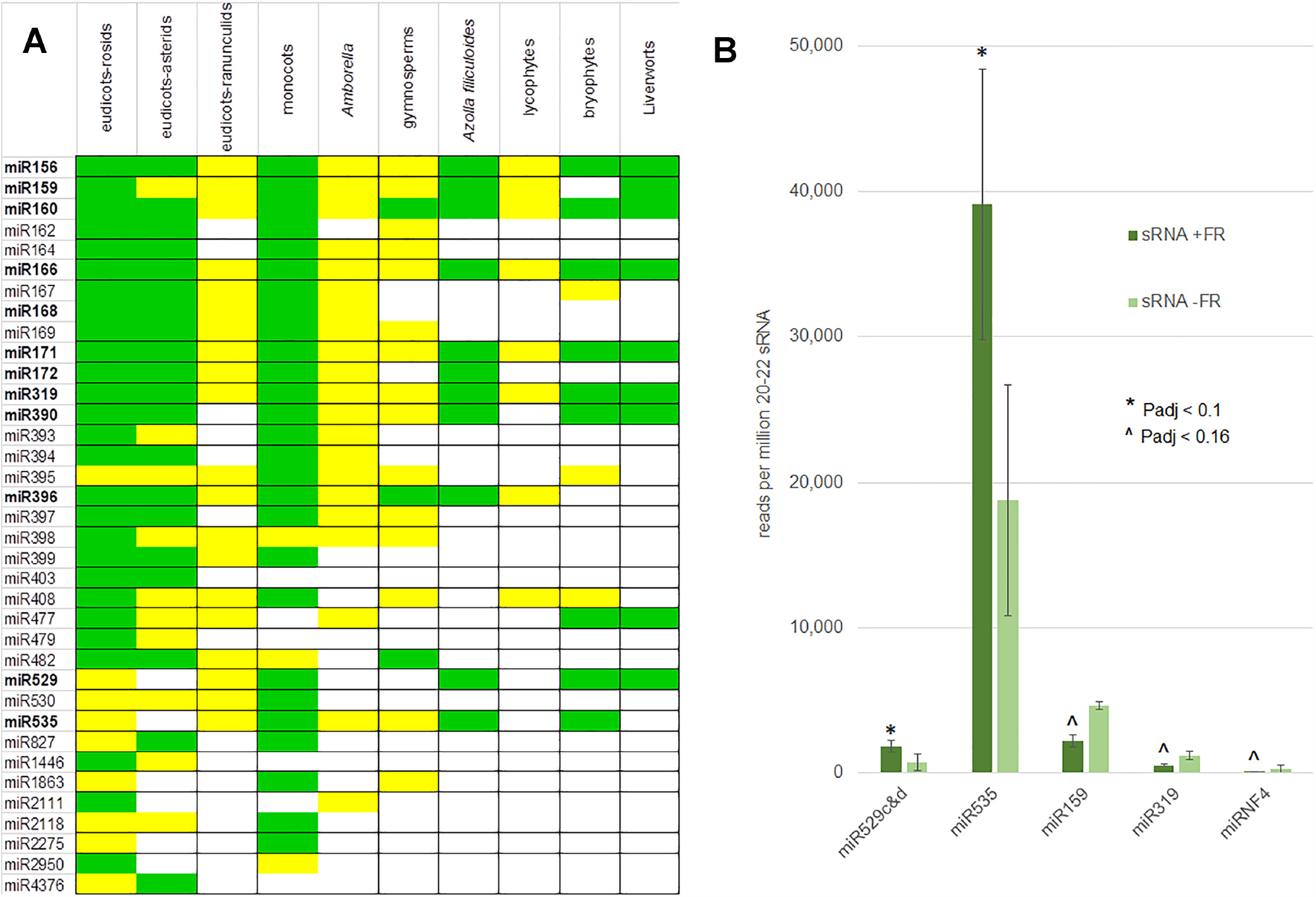
Conserved and far-red light responsive miRNA in the fern *A. filiculoides*. **(A)** Conserved miRNA in the *A*.*filiculoides* fern compared to other land plant lineages. miRNA from the fern were predicted computationally by miRDEEP-P2 using sRNA seq reads then curated manually, the remainder of the table was from Axtell & Meyers, 2018. Green, high confidence miRNA; yellow low confidence miRNA. **(B)** miRNA with altered steady state in response to far-red light. Shown are average read counts per million 20-22 nt reads in samples of sporophytes after one week on TL light with far-red LED (f) or without (t); raw read counts were submitted to DESeq2 for statistical analyses yielding P-adjusted values: * indicates Padj <0.1; ^ indicates Padj <0.16.

Differential expression analysis of 20-22 nt reads exhibited more dispersion than the dualRNAseq of long RNA, yet, it reliably identified sRNA with low Padj values when comparing ferns under TL with and without FR (Supplemental Figures 5D and 5E). Reads from miR529c&d and miR535 increased (Padj < 0.1) whilst those of miR159, miR319 and miRNF4 decreased (Padj <0.16) in sporophytes on TL with FR compared to TL light only (Figure 7B). We conclude that the miRNA156/miR172 known to reciprocally control flowering are not altered in the ferns undergoing transition to RD under FR.

### miR319/GAMYB is responsive in sporophytes induced by FR

miRNA targets were predicted combining psRNATarget (Dai *et al*., 2018) and TargetFinder (Srivastava *et al*., 2014) then annotated with Mercator (Lohse *et al*., 2014; Schwacke *et al*., 2019). Predictions were consistent with analyses in other land plant lineages (Supplemental Table 6): miR172 targeted AP2/TOE-like transcription factors and miR156, 529 and miR319 SPL-like proteins.

Transcripts of neither SPL nor AP2/TOE-like targets, however, were significantly (Padj < 0.1) altered in sporophytes comparing TL with and without FR; this could be due to poor Azfi_vs1 annotation and incomplete prediction of miR targets (Supplemental Table 6, F vs T Padj). Nevertheless, consistent with an increased miR529c&d the SPL from the Azfi_s0173.g055767 transcript is reduced (log2 0.74 fold) but with low significance (Padj of 0.356). In contrast, transcripts of two GAMYB (Azfi_s0004.g008455 and Azfi_s0021.g015882) increased up to 4 fold in sporophytes with decreased miR319 on TL with FR. The location of mir319 binding is shown for the GAMYB encoded in Azfi_s0004.g008455 (Supplemental Figure 8).

Phylogenetic analyses revealed that the GAMYB targeted by miR319 were on the same leaf as the GAMYB33 from Arabidopsis and were placed next to each other, probably because they arose from gene duplication (Figure 8). This is consistent with the whole genome duplication observed at the base of *Azolla* evolution (Li *et al*., 2018). We conclude, therefore, that miR319/GAMYB is responsive in sporophytes undergoing transition to RD caused by FR.

**Figure 8.**
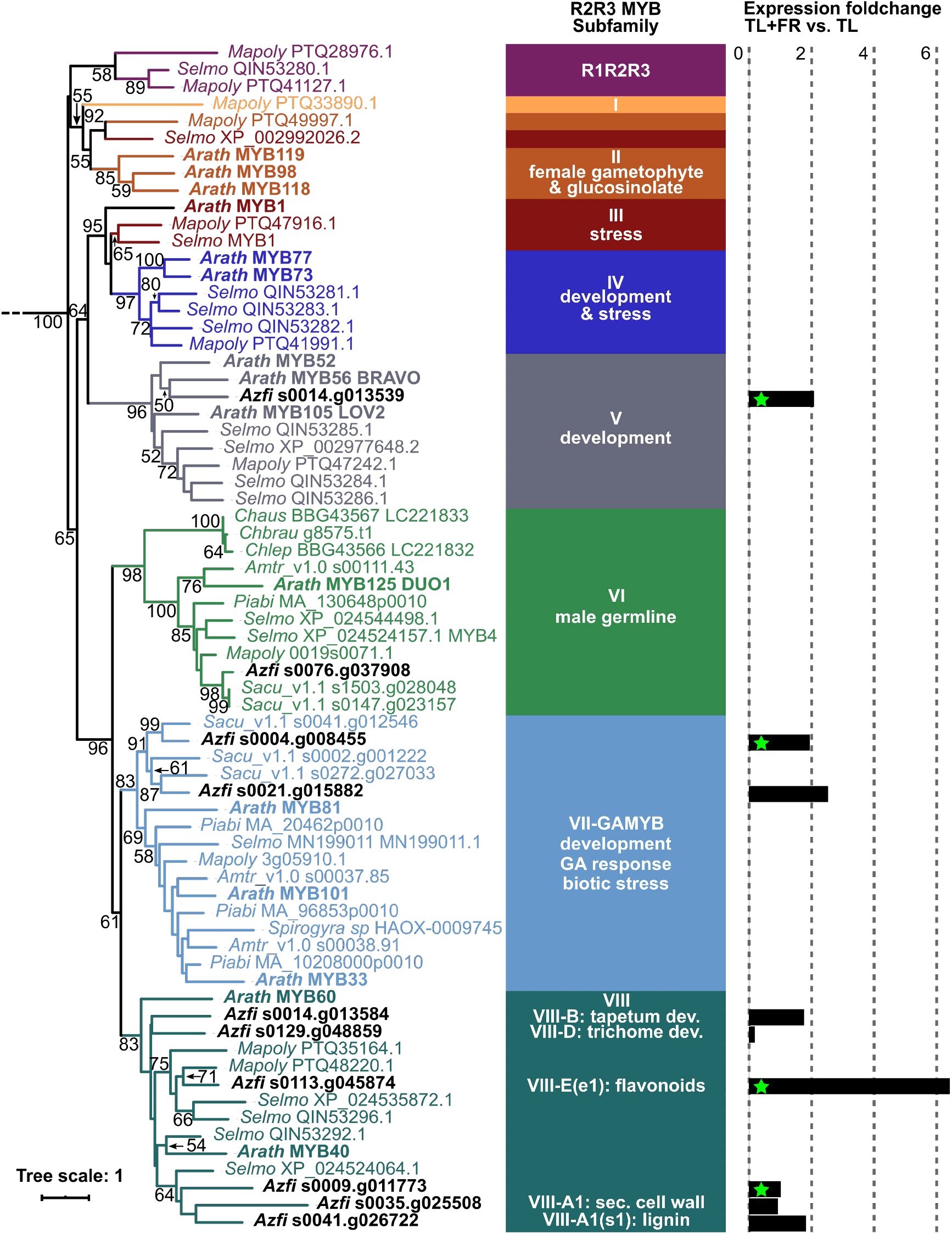
R2R3MYB phylogenetic analyses and fold change comparing sporophytes with and without FR LED. Sequences were extracted from the genome browsers of each species, aligned with MAFFT linsi or einsi, then trimmed with trimAL. Phylogenetic inferences were computed with IQtree and its internal model fitter; non-parametric bootstrap values are shown. *Chara autralis* (Ca), *Chara braunii* (Cb), *Chara leptospora* (Cl), *Marchantia polymorpha* (Mp), *Selaginella moellendorfii* (Sm), *Azolla filiculoides* (Azfi),), *Salvinia cuculata* (Sc), *Picea abies* (Pa), *Amborella trichopoda* (Atri), *Arabidopsis thaliana* (At). Sequence names are color coded after the R2R3 MYB subfamilies defined in Jiang & Rao (2020). Fold change in response to FR was calculated by DESeq2, green stars mark significant changes with Padj <0.1.

## DISCUSSION

### Red-dominated light suppresses formation of dissemination stages in both gametophyte and sporophyte-dominated lineages of plants and is, therefore, a convergent ecological strategy

Far-red light responses in *Azolla* ferns look alike a shade-avoidance syndrome but signal transduction pathways that mediate them in ferns remain largely uncharacterized (Inoue *et al*., 2017). Pathways causing shade response components are known to use alternative phytochromes and interacting factors (Possart *et al*., 2013; Xie *et al*., 2020).

Phytochromes in ferns radiated separately from those of seed plants (Li *et al*., 2015), hence, their function cannot be predicted using orthology with seed plants. Nevertheless, fern phytochromes have a similar structure and, as in the case of PHYB in Arabidopsis, may sense temperature as well as red/far-red light by thermal reversion of the Pfr to fr state (Legris *et al*., 2017; Klose *et al*., 2020): initiation of fern, liverwort and bryophyte sporangia was dependent on temperature and photoperiod (Labouriau, 1958, Benson-Evans, 1961; Nishihama *et al*., 2015). Moreover, thermal reversion of phytochromes from some cyanobacteria was described *in vitro* as early as 1997 (Yeh *et al*., 1997). Therefore, if far-red light induces RD in *A. filiculoides*, then differing temperature regimes combined with light quality changes may induce RD in the Anzali ferns. Anzali ferns were not *A. filiculoides* but related to species from Uruguay and Southern Brazil which did not cluster with, (Figure 2) and did not have the single-celled papillae of the traditional *A. caroliniana* species (Pereira *et al*., 2001; Supplemental Figure 3). Position of the Anzali accession was consistent with observations of distinct *Azolla* ferns in the Anzali lagoon (Farahpour-Haghani *et al*., 2017), and suggests that not only *A. filiculoides* may have been released as nitrogen biofertilizer.

Unlike *A. filiculoides*, Arabidopsis will transition to RD in the TL generally used for indoors cultivation. Most studies on the transition to RD, therefore, were done with red-light gated plants, i.e. conditions leading to artefactual peaking of CONSTANS protein accumulation and florigen (FT) expression (Song *et al*., 2018). Nevertheless, Arabidopsis as well as many other seed plants flower early when in ratios of red to far-red light approaching one, the ratio encountered in day light, compared to TL. Far-red light-dependent flowering is mediated by PHYB and D, eventually leading to *SOC1* expression in the meristem (Halliday *et al*., 1994; Aukerman *et al*., 1997; Lazaro *et al*., 2015). ft mutant Arabidopsis flowered earlier under incandescent light (Martinez-Zapater & Somerville, 1990) or in TL with far-red LED compared to in TL (Schluepmann *et al*., unpublished) suggesting that FT is not involved. Signaling was also reported via PHYB-regulated PIF4 protein complexes that bind the promoters of miRNA156/172 directly with interference from PHYA-regulated TF (Sánchez-Retuerta *et al*., 2018; Sun *et al*., 2018; Xie *et al*., 2020). In the present study, the highly induced MIKC^C^ turned out a paralogue of SOC1, and miR156/172 steady states were unaltered by the far-red triggered transition to RD in *A. filiculoides* (Figure 7B; Supplemental Table 6) suggesting that mechanisms in ferns and seed plant RD are not conserved.

Far-red induced sexual reproduction in the liverwort gametophytes *Marchantia polymorpha* (Chiyoda *et al*., 2008; Kubota *et al*., 2014; Yamaoka *et al*., 2018; Tsuzuki *et al*., 2019). The mechanism in these liverworts involved the MpmiR529c/MpSPL module, which was necessary for the development of reproductive branches. MpPHY and MpPIF were required to mediate the response (Inoue *et al*., 2019). Our analyses with *A. filiculoides* reveal induction of AzfimiR529 under far-red light but no significant change in SPL targets. Additionally, the specific MpmiR529c sequence involved in *M. polymorpha* was not detected in our ferns. The pathways thus differ in the two seed-free plants.

Given that gametophyte and sporophyte dominated lineages require far-red light to initiate formation of dissemination stages, we conclude that repression of RD under dominating red light in *Azolla* likely reflects a convergent ecological strategy: in open surfaces, where red-light is abundant, there is sufficient space to prolong vegetative development and no need for dispersal. Consistently, density of the *A. filiculoides* canopy had an additional impact on the number of sporocarps observed per gram plant (Figure 2F) that may stem from alternative cues such as volatile organic compounds released by neighbors (Vicherová *et al*., 2020). Taken together, results presented in this work signify that, for the first time, RD of the *Azolla* fern symbiosis is amenable to experimental enquiry.

### Phylogenetic position of MIKC^C^ TF responsive during *Azolla* RD suggests that control of flowering in seed plants originates from the diploid to haploid transition in the common ancestor of seed plants and ferns

The GAMYB TF clade arose before land plants evolved and before GA signaling (Aya *et al*., 2011; Bowman *et al*., 2017, Jiang & Rao, 2020, Figure 8). *Azolla* GAMYB induced under TL with far-red, therefore, likely function in RD by mediating cues perceived by the sporophyte into an ancestral gametophyte regulon. GAMYB from the moss *Physcomytrella patens* were required for the formation of antheridiophores by the gametophyte, and of normal spores by the sporophyte (Aya *et al*., 2011). The GAMYB regulon did not play a role in the developmental transition of lycophyte sporophytes to formation of sporogenic structures.

AtGAMYB are known to be regulated by miR159 in Arabidopsis and by the related and more ancestral miR319 in seed-free plant lineages (Achard *et al*., 2004; Palatnik *et al*., 2007). The AtGAMYB/miR159 is part of the GA pathway promoting flowering in Arabidopsis under short days (Millar *et al*., 2019). Specifically, AtMYB33 was shown to directly act on the promotors of both miR156 and AtSPL9 (Guo *et al*., 2017). Far-red light repressed miR159 and miR319, with increased AzfiGAMYB, in *Azolla* were thus consistent with wiring in Arabidopsis; furthering their analysis in homosporous ferns with free gametophytic stages will reveal whether they are important for the diploid to haploid transition.

Control over sexual reproductive transition as it switched from the gametophyte, in gametophyte dominated seed-free plants, to the diploid sporophyte in seed plants, resulted from the evolution of inflorescence and floral meristem MIKC^C^ regulons. The MIKC^C^ clade of TF that specify both the transition to RD in seed plant sporophytes, such as AtSOC1 and AtFLC, and the A,C, D and E homeotic functions are peculiar in that the TF radiated in each megaphyll plant lineage separately (Leebens-Mack *et al*., 2019; Supplemental Figure 4). The TF work as hetero-tetramers and may have evolved from an ancient tandem gene duplication through subsequent polyploidy events (Zhao *et al*., 2017). Sepals are organs formed only in angiosperms. Consistently, TF with A functions specifying sepals occurred solely in angiosperms. MIKC^C^ specifying organs that contain the reproductive structures in angiosperms (C, D and E homeotic genes), had direct homologues in gymnosperms, but not in ferns. Instead, ferns radiated a sister clade to the clade containing AtSOC1, and A, C, D and E MIKC^C^. This clade contained the *Azolla* MIKC^C^ responsive to far-red light and the closely related *Ceratopteris* CMADS1. *In situ* hybridizations and northern blot analyses reveal that *CMADS1* transcripts accumulate at high levels in the sporangia initials (Hasebe *et al*., 1998). At the base of the radiating clade of modern MIKC^C^ that specify modern organs of the sporophytes controlling the gametophyte development within, is the LAMB1 protein expressed specifically in sporogenic structures of lycophytes (Leebens-Mack *et al*., 2019; Svensson *et al*., 2000; and Supplemental Figure 4). Combined phylogenetic and RNAseq analyses, therefore, suggest that the MIKC^C^ regulons controlling flowering originate from the diploid to haploid phase transition of the common ancestor to ferns and seed plants, not from regulons controlling sexual reproduction of the haploid phase.

### Dual RNA sequencing to study coordinate development in assemblages of pro- and eukaryotic organisms

The dual RNAseq method could be applied more generally to study bacteria/plant assemblages particularly common in the seed-free plant lineages, representing an important base of the “tangled tree” of plant lineage evolution (Quammen D. 2018). For *Azolla* where co-evolution of *N. azollae* and fern host genomes demonstrated that fitness selection occurs at the level of the metagenome (Li *et al*., 2018), it delivered foundational data and opens the way to studying the coordinate development of host and obligate symbiont. Efficient removal of the rRNA was key to the success of reading long RNA. Fern chloroplast rRNA removal could be improved still, using specific DNA probes integrated during synthesis of the stranded sequencing libraries. Transcripts from the fern’s mitochondrial genome are likely in the data but we have yet to assemble the *A. filiculoides* mitochondrial genome to confirm this.

*N. azollae* extracts did not contain photosynthesis pigments absorbing/fluorescing in the far-red spectrum and the *N. azollae* genome did not contain psbA4/Chlf-S like proteins (data not shown). Yet genes encoding phytochromes and cyanobacteriochromes capable of sensing far-red light as other filamentous cyanobacteria are present (Wiltbank & Kehoe, 2019). The RNA profiling did not reveal whether the *N. azollae* trigger fern RD since roles of the far-red responsive transcripts in *N. azollae*, particularly those encoding the transporters, have yet to be studied. An approach could be to study these in free living filamentous cyanobacteria amenable to genetic manipulation. Manipulation of the symbiont would, however, be preferred. Tri-parental mating protocols may be effective on *N. azollae in situ* if combined with high-frequency RNA-guided integration with Cas12d enzyme complexes (Strecker *et al*., 2019). *N. azollae* Cas genes were found to be mostly pseudogenes and CRISPR arrays appeared missing (Cai *et al*., 2013), suggesting that incoming cargo plasmid may not be destroyed by Cas complexes. Nevertheless, sRNA profiles from Figure 6 show discrete accumulation of 31 and 42 b sRNA of very low complexity and a functional Cas6 splicing CRISPR precursor RNA was induced on TL with far-red LED (Supplemental Table 4, *N. azollae*). CRISPR arrays may not be recognized if they contained few repeats due to the sheltered life-style inside the fern. Rendering *A. filiculoides* and *N. azollae* amenable to genetic manipulation will be crucial to provide evidence for hypotheses generated by transcript profiling in the future.

## MATERIALS AND METHODS

### Plant Materials, growth conditions and crosses

Ferns were collected from differing locations in the Netherlands, grown in liquid medium at a constant temperature of 22°C under long days (16h light) as described earlier (Brouwer *et al*., 2017). Ferns from Spain and Iran were processed to DNA on site. GPS coordinates from the collection sites of the strains used in phylogenetic analyses are in Supplemental Table S1. For sporocarp induction, ferns were first raised under 16 h tube light (TL) periods, then transferred for induction under 16 h TL with far-red Light Emitting Diodes - LED (Phillips GrowLED peaking at 735 nm, Supplemental Figures 1A and 1B). Photosynthetic Photon Flux Density varied in the range 100-120 μmol m^-2^ s^-1^. Sporophytes were generally kept at densities that covered the surface as this inhibits growth of (blue) algae. When testing sporocarp induction sporophytes were kept at set densities (with fresh weight resets once per week, Supplemental Figures 1C, 1D and 1E) and set red to far-red light ratios (Supplemental Table **2**). Phenotypic change in mature sporophytes were photographed using a Nikon D300s with an AF Micro-Nikkor 60mm f/2.8D objective or bellows on a rail to generate image reconstructions with Helicon focus software. Megasporocarps were collected manually using tweezers, so were microsporocarps. To initiate crosses microsporocarps were Dounce-homogenized and massulae thus released added to megasporocarps in distilled water (3-6 ml, in 3 cm diameter petri dish), shaking so as to obtain clumps. The crosses were kept at room temperature (about 18°C) under 16 h light conditions under TL at about 100 μmol m^-2^ s^-1^. As sporelings emerged they were transferred to liquid medium (Brouwer *et al*., 2017).

**RNA extraction and quantitative RT-PCR** were as in (Brouwer *et al*., 2017) and are detailed in Supplemental Methods S1.

**Phylogenetic analyses** are described in details in Supplemental Methods. Chloroplastic marker regions were as in Supplemental Table 3. The MIKC^C^ tree was subsampled from 1 KP (Leebens-Mack *et al*., 2019); the R2R3 MYB tree used data from respective genome repositories, the *Azolla* genome (Li *et al*., 2018) and Jiang & Rao, 2020.

**Reference genomes** used to align Dual RNA sequences onto the *A. filiculoides* Galgenwaard strain are detailed in Supplemental Methods.

### Dual-RNA sequencing and data processing

*A. filiculoides* sporophytes were from the sequenced accession Galgenwaard (Li *et al*., 2018). The ferns were maintained on liquid medium without nitrogen in long-day TL, then transferred to TL with and without far-red LED for 7 days, on fresh liquid medium without nitrogen; harvest was 2 h into the light period by snap freezing. All samples were grown separately and harvested as triplicate biological replicates. Total RNA was extracted using the Spectrum Plant Total RNA (Sigma-Aldrich) using protocol B, then DNAse treated as described in (Brouwer *et al*., 2017), and cleaned using the RNeasy MinElute Cleanup Kit (Qiagen). rRNA depletions were carried out as per protocol using Ribo-Zero rRNA Removal Kit (Plant Leaf, Illumina), and the RNA was then cleaned again using the RNeasy MinElute Cleanup Kit. Stranded libraries were then synthesized using the Ovation Complete Prokaryotic RNA-Seq Library Systems kit (Nugen). The libraries thus obtained were characterized and quantified before sequencing using a single lane Hiseq4000 and 50 cycles PE reading (Illumina).

Reads obtained were de-multiplexed then trimmed and quality filtered with trimmomatic (Bolger *et al*., 2014), yielding 13-60 million quality PE reads per sample. Alignment of the reads to the genome assembly and GFF vs1 of *A. filiculoides* (Azfi_vs1; Li *et al*., 2018) was with STAR-aligner default settings (Dobin *et al*., 2013) and was combined with the STAR read count feature (PE read counts corresponding to the HTseq “Union” settings, (Anders *et al*., 2015)). Read count tables were fed into DESeq2 (Love *et al*., 2013) for analysis of differentially accumulating transcripts.

STAR was also used to align reads to the *N. azollae* and the chloroplast genomes, and read counts then extracted using STAR default parameters or VERSE (Zhu *et al*., 2016) with HTSeq setting “Union” comparing single and paired End read counts on the polycistronic RNA. The three approaches yielded similarly ranked genes but the Padj statistics of differential expression in count tables from the forced single read counts were poor: only 20 *N. azollae* genes had P-adjusted values smaller than 0.1 compared to up to 67 with the PE-approaches, we therefore present the PE-approach using STAR read counts only.

STAR alignment to concatenated genomes of the *Azolla* symbiosis using default settings yielded a large proportion of multi-mappers which interfered with PE read counting leading to losses of as much as 30% of the reads. The strategy would reduce force mapping but is not necessary to extract differentially accumulating transcripts. We, however, used alignments allowing no mismatches to concatenated genomes when mapping the sRNA reads below, then recovering reads aligning to each genome using Samtools fasta from the indexed .bam files.

### Small RNA data sequencing and data processing

Ferns were maintained on liquid medium without nitrogen with 16 h TL, then transferred to TL with and without far-red LED for 7 days as described under Dual-RNA sequencing. In addition, sporophytes were grown on medium with or without 2 mM NH_4_NO_3_; ferns without cyanobacteria were grown on medium with 2 mM NH_4_NO_3_ in TL with far-red LED. All sporophytes were harvested 7 d into the treatment, 2 h into the light period snap frozen. Samples were harvested from triplicate growth experiments and thus represent triplicate biological replicates. Small RNA was extracted as per protocol using the mirPremier microRNA Isolation Kit (Sigma-Aldrich), then DNAse treated (Brouwer *et al*., 2017), then cleaned using the RNeasy MinElute Cleanup Kit (Qiagen) protocol for small RNA. Libraries of the sRNA were then generated using the SeqMatic TailorMix miRNA Sample Preparation 12-reaction Kit (Version 2) as per protocol. Libraries were characterized using an Agilent Technologies 2100 Bioanalyzer and quantified before sequencing using NextSeq500 and 1 x 75 b read chemistry (Illumina).

Reads thus obtained were de-multiplexed, trimmed and quality filtered with trimmomatic, then aligned with STAR (Dobin *et al*., 2013) (set at maximal stringency and to output Bam files) to the concatenated fasta files of the genomes Azfi_vs1, its chloroplast, and *N. azollae* 0708. The resulting bam files were then split for each genome, and Samtools fasta (Li *et al*., 2009) used to extract reads aligning to each genome. fastx collapser (http://hannonlab.cshl.edu/fastx_toolkit) was then used to extract read counts for each identical sequence, with the sequence defining the name in the lists obtained. A data matrix was generated in bash, then exported to Excel or Rstudio for further analyses.

To determine differential expression, the lists obtained were restricted to sRNA with at least 10 reads per sample joined in bash, and the tables thus obtained, first restricted to containing only reads of 20-22 nt sRNA (to minimize gel excision bias during sequencing library preparation), then exported to R (Booth *et al*., 2018). DESeq2 was then used to display variation and dispersion in the data and identify differentially accumulating sRNA using Padj values (Love *et al*., 2013).

### miRNA discovery

The miRNA loci in *A. filiculoides* were analyzed as in Supplemental Figure **2**. Pre-miRNA were predicted with miRDEEP-P2 (Kuang *et al*., 2019); multi-mappers min. 100 and max. loop length 500) then extracted and subjected to secondary structure prediction using RandFold default parameters (Bonnet *et al*., 2004; Gruber *et al*., 2008). miRNA candidates containing less than four mismatches within the hairpin structure were viewed in IGV along with RNA expression data aligned to Azfi_vs1. Predicted miRNAs that were expressed in *A. filiculoides* were then compared in miRbase (Kozomara *et al*., 2019). Prediction of miRNA targets in Azfi_vs1 was using the intersect of both psRNATarget (Dai *et al*., 2018) and TargetFinder (Srivastava *et al*., 2014) with a cutoff of 5.0. Predicted targets were linked to the closest homolog using the Mercator annotation (Lohse *et al*., 2014) of Azfi_vs1 predicted proteins.

### Data sharing

The data that support the findings of this study are openly available in [repository name] at [URL], reference number [reference number]. Data will be uploaded upon acceptance of the manuscript and will include 1) read files for the dual RNA seq as well as small RNA seq on ENA 2) phylogeny files on Github.

## Supporting information

Supporting Information Methods, Figures and compact Tables

## Supplemental Data

### Supplemental Methods

RNA extractions & quantitative RT-PCR, phylogenetic analyses, and metagenome assemblies of *N*.*azollae* genomes from different *Azolla* species.

**Supplemental Table 1**. Location and coordinates of *Azolla* collection sites for strains used in the phylogenetic analyses.

**Supplemental Table 2**. Red to far-red light intensity ratios and culture densities when testing induction ofsporulation by far-red light.

**Supplemental Table 3**. Sequence accessions of the fern accessions from the phylogenetic trees in Figure 2A and 2B.

(separate Excell file with sheets for the Trn L-F and Trn G-R).

**Supplemental Table 4**. Differentially accumulating transcripts in sporophytes on tube light with and without

far-red LED. (separate Excel file).

**Supplemental Table 5**. Read counts for transcripts of *N. azollae*. (separate Excell file).

**Supplemental Table 6**. miRNA and target loci transcript abundance in sporophytes in response to FR light.

**Supplemental Figure 1**. Light quality and fern densities when testing induction of sporulation. **Supplemental Figure 2**. Flow chart to discover conserved and novel miRNA in *A. filiculoides*. **Supplemental Figure 3**. Taxonomy of the *Azolla* strains in this study.

**Supplemental Figure 4**. Phylogenetic analysis of the *A. filiculoides* MIKCC responsive to FR.

**Supplemental Figure 5** sRNA sequencing and mapping statistics.

**Supplemental Figure 6**. MiR172 target sites in AP2/TOE1 transcription factors (eaAP2 lineage) comparing *A*.*filiculoides* and seed plants.

**Supplemental Figure 7**. The miRNA156α locus in A. filiculoides.

**Supplemental Figure 8**. The AzfiGAMYB locus of A.filiculoides targeted by miRNA319.

## ACKNOWLEDGEMENTS

NWO-ALW EPS grant (ALWGS.2016.020) supported LWD, NWO-TTW grant (Project 16294) supported EG. We are grateful to Bruno Huettel for advice on dual RNA sequencing library preparation methods. We thank Bas Dutilh and Berend Snel for access to the Utrecht University bioinformatic computing resources.

## AUTHOR CONTRIBUTIONS

Fern induction experiments were done by BST and PB, fern crosses and comparisons were done by EG, BST and unnamed graduate students. Images of shoot apices were generated by VB, phylogenetic analyses and miR discovery were carried out by LWD and NR, sRNA and dual RNA profiling was done by HS. HS conceived the manuscript with all authors contributing to editing it.

