## Supporting Information Methods, Figures and compact Tables for "Control of the *Azolla* symbiosis sexual reproduction: ferns to shed light on the origin of floral regulation?"

The following Supplemental Data is available for this article:

**Supplemental Methods.** RNA extractions & quantitative RT-PCR, phylogenetic analyses, and metagenome assemblies of *N.azollae* genomes from different *Azolla* species. **Supplemental Table 1.** Location and coordinates of *Azolla* collection sites for strains used in the phylogenetic analyses.

**Supplemental Table 2.** Red to far-red light intensity ratios and culture densities when testing induction of sporulation by far-red light.

**Supplemental Table 3.** Sequence accessions of the fern accessions from the phylogenetic trees in Figures 2A and 2B. (separate Excell file with sheets for the Trn L-F and Trn G-R).

**Supplemental Table 4.** Differentially accumulating transcripts in sporophytes on tube light with and without far-red LED. (separate Excel file).

**Supplemental Table 5.** Read counts for transcripts of *N. azollae*. (separate Excell file).

**Supplemental Table 6.** miRNA and target loci transcript abundance in sporophytes in response to FR light.

**Supplemental Figure 1.** Light quality and fern densities when testing induction of sporulation.

**Supplemental Figure 2.** Flow chart to discover conser ved and novel miRNA in *A. filiculoides*.

**Supplemental Figure 3.** Taxonomy of the *Azolla* strains in this study.

**Supplemental Figure 4.** Phylogenetic analysis of the *A. filiculoides* $MIKC^C$ responsive to FR.

**Supplemental Figure 5.** sRNA sequencing and mapping statistics.

**Supplemental Figure 6.** MiR172 target sites in AP2/TOE1 transcription factors (eaAP2 lineage) comparing *A. filiculoides* and seed plants.

**Supplemental Figure 7.** The miRNA156α locus in *A. filiculoides*.

**Supplemental Figure 8.** The *AzfiGAMYB* locus of *A.filiculoides* targeted by miRNA319.

### Supplemental Methods

#### RNA extraction from sporophytes and quantitative RT-PCR

RNA was extracted, DNAse treated then reverse transcribed (Brouwer *et al.*, 214). Primers for qRT-PCR were for the references *AfTUBULIN* (AfTUBF: CCTCCGAAAACTCTCCTTCC; AfTUBR: GGGGGTGATCTAGCCAAAGT) and *AfADENINE PHOSPHORIBOSYLTRANSFERASE* (AfAPTF: TAGAGATGCATGTGGGTGCAGT; AfAPTR: AAAAGCGGTTTACCACCCAGTT), and for *AfSOC1* (AfSOCF: ATGGGATCGTAAGGCTTCAAAA; AfSOCR: AGCAGAGCACACAGGTCTCAAC). qRTPCR amplifications were from RNA of three biological replicate growth tubs, significance was assessed by *t*-test with $P < .5$.

#### Phylogenetic analyses

Chloroplastic marker regions. DNA was extracted using the extraction kit (EZNA SP Plant DNA), PCR amplifications were using *pfu* and *taq* polymerase mixed (1:4 Units) with *pfu*-bufffer containing $MgSO_4$ (2 mM) and *taq*-cycling conditions (37 cycles of 3 s 94°C denaturation, 3 s 5-58°C annealing, 1 min 72°C amplification, then 5 min 72 °C) and using primers trnG1F (GCGGGTATGGTTTAGTGGTAA), trnR22R (CTATCCATTAGACGATGGACG), trnLC (CGGAATGGTAGACGCTACG) and trnLF (ACTTGAACTGGTGACACGAG). Amplicons were purified (EZNA Cycle pure kit) then submitted for sequencing (Macrogen). Sequences were reconstructed then added to a fasta file containing the cognate sequences from Azolla species sequenced previously (Madeira *et al.*, 216; Dijkhuizen *et al.*, 218). Sequences were aligned on MEGA 7 (Kumar *et al.*, 216) using Clustal W (Thompson *et al.*, 1994). Gap opening and gap extension of 1 and 3, respectively, were used for trnL-trnF and 1 and 4, respectively, were used for trnG-trnR. Maximum likelihood phylogenetic trees were calculated with alignment lengths of 92 bp and 852 bp for trnL-trnF and trnG-trnR, respectively. The trnL-trnF tree was constructed using the Tamura 3-parameter plus Gamma (T92 + I) model and the trnG-trnR tree using the Tamura 3-parameter plus Invariant (T92 + I) model. The gaps/missing data treatment was chosen as "partial deletion", branch swap filter as "very weak" and ML Heuristic method as "Subtree-Pruning-Regrafting - Extensive (SPR level 5)". The reliability of branches was analyzed by using bootstrapping of 1 replicates; trees obtained were visualized with iTOL (Letunic & Bork, 216).

Sequence logo of ITS1 intergenic regions. The logo from the *A. filiculoides* ITS was generated using WebLogo3 (Crooks *et al.*, 24) after extracting 516 sequences from the Azfi vs 1 genome with *in silico* PCR (iPCRess, Slater & Birney, 25) and the published reference sequence for *A. filiculoides* (Li *et al.*, 218). ITS1 sequences other Azolla species (Dijkhuizen *et al.*, 218) were extracted by alignment to the *A. filiculoides* reference sequence and the Bam file obtained then used to generate the sequence logos of ITS regions from other *Azolla* species.

MIKC$^C$ and R2R3MYB phylogenetic tree. The automatically generated annotation of Azfi vs1 was first corrected in IGV using reads from the dual RNAsequencing experiment. Azolla fern sequences were then merged to those extracted from the genome browsers of each species and aligned with MAFFT linsi or einsi (Katoh *et al.*, 219), then trimmed with trimAL (Capella-Gutirrez et al., 29). Phylogenetic inferences were computed with IQTREE (Nguyen *et al.*, 215) and its internal model fitter (Trifinopoulos *et al.*, 2016). The resulting maximum likelihood (ML) tree was visualized in iTOL (Letunic & Bork, 2019) with minimum bootstrap support of 50% and sequences color-coded based on their clade assignment (R2R3MYB) or on phylogenetic placement ($MIKC^C$).

**Reference genomes used in Dual RNA sequencing**

Assembly and annotation of the *A.filiculoides* accession Galgenwaard chloroplast and nuclear genomes were from (Li *et al.*, 2018). Metagenome Assembled Genomes (MAGs) from *N. azollae* in the different *Azolla* species were computed using the shotgun sequencing data of the accessions in Dijkhuizen *et al.*, 2018: trimmed and quality assessed reads were filtered (BWA aligner,(Li & Durbin, 2009)) against the fern and chloroplast genomes then assembled into contiguous sequences (contigs) using SPAdes in metagenome mode (Nurk *et al.*, 2017), and contigs assigned Streptophyta taxonomy with CAT (Meijenfeldt *et al.*, 2019) added to the nucleus and chloroplast filter for another round of read filtering. Reads obtained were then assembled with SPAdes again, then contigs binned according to k-mer, GC and vertical coverage of the reads on the contigs. Bins were further interactively polished in Anvi'o (Eren *et al.*, 2011) Average Nucleotide Identity (ANI) was calculated with the dRep implementation of nucmer using the ANImf preset (Kurtz *et al.*, 2004; Olm *et al.*, 2017).

**Supplemental Table 1. Location and coordinates of *Azolla* collection sites for strains used in the phylogenetic analyses.**

| Geographic location | Coordinates |
|---|---|
| Galgenwaard, Utrecht, Netherlands | 52°4’35.73”N, 5°8’59.05”E |
| Nieuwerbrug, Netherlands | 52°04'45.5"N 4°48'28.1"E |
| Hoogwoud, Netherlands | 52°43'14.7"N 4°56'25.6"E |
| Kromme, Netherlands | 52°05'02.4"N 5°09'32.1"E |
| Den Bosch, Netherlands | 51°41'12.5"N 5°20'17.0"E |
| Nijmegen, Netherlands | 51° 49' 25.1'' N 5° 52' 9.501'' E |
| Gran Canaria, Spain | 28°04'00.9"N 15°27'38.9"W |
| Anzali lagoon, Iran | 37° 28' 8.007'' N 49° 21' 13.208''E |
| *Azolla mexicana* Schltdl. & Cham. ex Kunze | IRRI accession ME2001; originally from USA, California, Graylodge, collected by D. Rains in 1978 |
| *Azolla microphylla* Kaulf. | IRRI accession MI4021; originally from Ecuador, Galapagos, Santa Cruz Island; collected by T. Lumpkin in 1982 |
| *Azolla nilotica* Mett. | IRRI accession NI5001; originally from Sudan, Kosti; collected by T. Lumpkin in 1982 |
| *Azolla caroliniana* Willd. | IRRI accession CA3017; originally from Brazil, Rio Grande Sul; collected by I. Watanabe in 1987 |
| *Azolla caroliniana* | IRRI accession CA3004; originally from Uruguay, Treinta y tres; collected by D. Rains in 1982 |
| *Azolla rubra* R. Br. | IRRI accession RU6502; originally from Australia, Victoria, collected in 1985 |

**Supplemental Table 2. Red to far red light intensity ratios and culture densities when testing induction of sporulation by far-red light. ± refers to the variation from three replicate cultures maintained over a period of 6 weeks.**

| Light condition | Red:far-red ratio | Density range of the fern culture [DW g $m^{-2}$] | | |
|---|---|---|---|---|
| | | +/- 50 | +/-140 | +/- 240 |
| TL | 19,13 | 45 ± 23 | 139 ± 19 | 242 ± 18 |
| FR-1 | 1,16 ± 0,18 | 49 ± 26 | 142 ± 22 | 242 ± 15 |
| FR-2 | 0,63 ± 0,09 | 48 ± 25 | 137 ± 18 | 239 ± 17 |
| FR-3 | 0,50 ± 0,06 | 49 ± 26 | 140 ± 21 | 242 ± 19 |
| FR-4* | 0,31 ± 0,02 | 40 ± 16 | 126 ± 8 | 221 ± 32 |

*To achieve the ratio in FR-4, the total PAR was reduced by half.

**Supplemental Table 3. Trn L-F and Trn G-R Sequence accessions used for the phylogenetic trees in Figure 2A and 2B.** (separate Excel file with sheets for the Trn L-F and Trn G-R).

**Supplemental Table 4. Differentially accumulating transcripts in sporophytes on tube light with and without far-red LED.** (separate Excel file with sheets for *N. azollae*, *A. filiculoides* chloroplast and *A.filiculoides* nucleus).

**Supplemental Table 5. Read counts for gene features of *N. azollae* inside *A. filiculoides* sporophytes grown under TL without and with far-red LED.** (separate Excel file).

**Supplemental Table 6. miRNA and target loci transcript abundance in sporophytes on TL with versus without far-red LED.**

MiR, miRNA; target locus, MiR target locus; DESeq2 base mean, corrected mean expression computed by DESeq2,; log2-fold, log2-fold change comparing the three replicates of ferns exposed to far-red to those without on tube light only (F vs T); Padj, adjusted P-value according to DESeq2 comparing F vs T; Mercator, annotation of the target locus as predicted by Mercator. Targets with a Padj <0.122 are in bold; those with Padj <0.4.2 are in grey.

| | **Target locus** | **DESeq2** | **F vsT** | **F vsT** | |
|---|---|---|---|---|---|
| **MiR** | | base mean | log2-fold | Padj | Mercator |
| **miR156a,b** | Azfi_s0173.g055767 | 313 | 0.743 | 0.355805 | SPL-like |
| | Azfi_s0093.g043231 | 308 | 0.775 | 0.419548 | SPL-like |
| | Azfi_s0048.g030445 | 275 | 0.543 | 0.91417 | SPL-like |
| | Azfi_s0052.g031491 | 273 | 0.350 | 0.95265 | SPL-like |
| | Azfi_s0211.g058143 | 37 | 0.251 | 0.993074 | SPL-like |
| | Azfi_s0068.g036283 | 22 | -0.069 | 0.996389 | SPL-like |
| | Azfi_s0213.g058306 | 522 | -0.020 | 0.997365 | SPL-like |
| **miR529a,b,e,f** | Azfi_s0048.g030445 | 275 | 0.543 | 0.91417 | SPL-like |
| | Azfi_s0052.g031491 | 273 | 0.350 | 0.95265 | SPL-like |
| | Azfi_s0211.g058143 | 37 | 0.251 | 0.993074 | SPL-like |
| | Azfi_s0213.g058306 | 522 | -0.020 | 0.997365 | SPL-like |
| **miR529c,d** | Azfi_s0173.g055767 | 313 | 0.743 | 0.355805 | SPL-like |
| | Azfi_s0052.g031491 | 273 | 0.350 | 0.95265 | SPL-like |
| | Azfi_s0211.g058143 | 37 | 0.251 | 0.993074 | SPL-like |
| | Azfi_s0068.g036283 | 22 | -0.069 | 0.996389 | SPL-like |
| **miR535** | Azfi_s0093.g043231 | 308 | 0.775 | 0.419548 | SPL-like |
| | Azfi_s0048.g030445 | 275 | 0.543 | 0.91417 | SPL-like |
| | Azfi_s0213.g058306 | 522 | -0.020 | 0.997365 | SPL-like |
| **miR319** | **Azfi_s0004.g008455** | **103** | **1.930** | **0.002829** | **R2R3-GAMYB** |
| | **Azfi_s0021.g015882** | **25** | **2.514** | **0.121226** | **R2R3-GAMYB** |
| | Azfi_s0460.g072042 | 7 | 1.699 | 0.932033 | R2R3-GAMYB |
| | Azfi_s0138.g051134 | 21 | 0.433 | 0.977615 | R2R3-MYB |
| | Azfi_s0041.g026717 | 44 | -0.255 | 0.984461 | R2R3-MYB |
| **miR172a,d** | Azfi_s0178.g056175 | 1,319 | 0.545 | 0.792255 | AP2-like |
| | Azfi_s0496.g073599 | 636 | -0.281 | 0.963591 | AP2-like |
| **miR172b,c** | Azfi_s0178.g056175 | 1,319 | 0.545 | 0.792255 | AP2-like |
| | Azfi_s0496.g073599 | 636 | -0.281 | 0.963591 | AP2-like |
| **miR160a** | Azfi_s0002.g001367 | 52 | 0.624 | 0.859618 | ARF |
| | Azfi_s0019.g015120 | 243 | -0.466 | 0.900243 | ARF |
| | Azfi_s0099.g044147 | 633 | -0.467 | 0.914563 | ARF |
| | Azfi_s0013.g013413 | 377 | -0.507 | 0.940884 | ARF |
| | Azfi_s0697.g082598 | 168 | 0.054 | 0.996389 | ARF |
| **miR160c,d** | Azfi_s0002.g001367 | 52 | 0.624 | 0.859618 | ARF |
| | Azfi_s0019.g015120 | 243 | -0.466 | 0.900243 | ARF |
| | Azfi_s0099.g044147 | 633 | -0.467 | 0.914563 | ARF |
| | Azfi_s0013.g013413 | 377 | -0.507 | 0.940884 | ARF |
| | Azfi_s0697.g082598 | 168 | 0.054 | 0.996389 | ARF |
| **miR171a** | Azfi_s0019.g015002 | 1,193 | -0.225 | 0.975528 | GRAS |
| | Azfi_s0032.g024755 | 311 | -0.208 | 0.977615 | GRAS |
| **miR171b** | Azfi_s0019.g015002 | 1,193 | -0.225 | 0.975528 | GRAS |

**Supplemental Figure 1. Light quality and fern densities when testing induction of sporulation**.
(**A)** Spectrum of the light under tube light (TL);
(**B)** Spectrum of the light under TL with far-red LED.
Fern densities over 6 weeks of sporocarp induction: cultures refreshed weekly at 41.7 (**C)**, 128.8 **(D)** or 208.3 **(E)** g DW $m^{-2}$. Cultures were weighed and densities re-set at the beginning of each week. Data points are averages of three replicate cultures with standard deviations.
**(F)** Proportion of mega to microsporocarps over a 9 week induction experiment;
**(G)** *AzfiSOC1* mRNA accumulation detected by qRT-PCR in sporophytes grown under TL (no FR) and TL with far-red LED (Far Red) for 7 weeks.

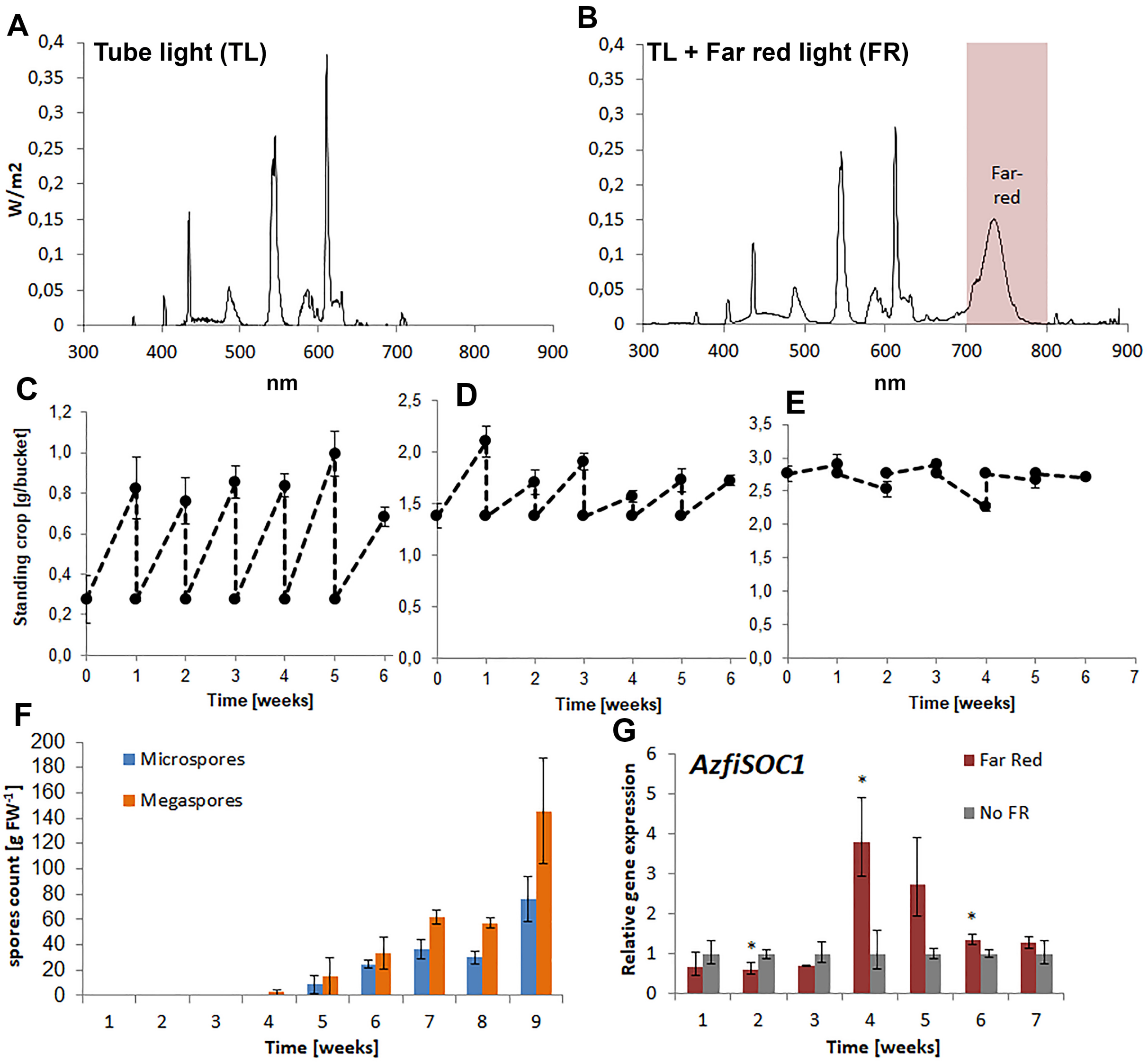

**Supplemental Figure 2. Flow chart of the discovery of conserved and novel miRNA in *A. filiculoides*.** On the one hand, the miRNA predicted by You et al., 2014 for *A. caroliniana* were used for a Blast search of the *A. filiculoides* genome and hits viewed and further analyzed using IGV (Thorvaldsdóttir et al., 2013). On the other hand, sRNA seq reads were quality filtered, then collapsed for identical read sequence then only 20-22 nt reads retained for further analyses, the reads were sorted then submitted for analysis with miRDEEP2 and mirDEEP-P2 (Kuang et al., 2018); resulting candidates were verified manually for fold potential in Vienna Fold (Gruber et al., 2008) and expression using IGV, then compared with existing miRNA in miRbase vs 22.1 (Kozomara et al., 2014). Finally, candidate targets were explored using the intersection of results from Targetfinder (Fahlgren & Carrington, 2010) and psRNATarget (Dai *et al.*, 2018).

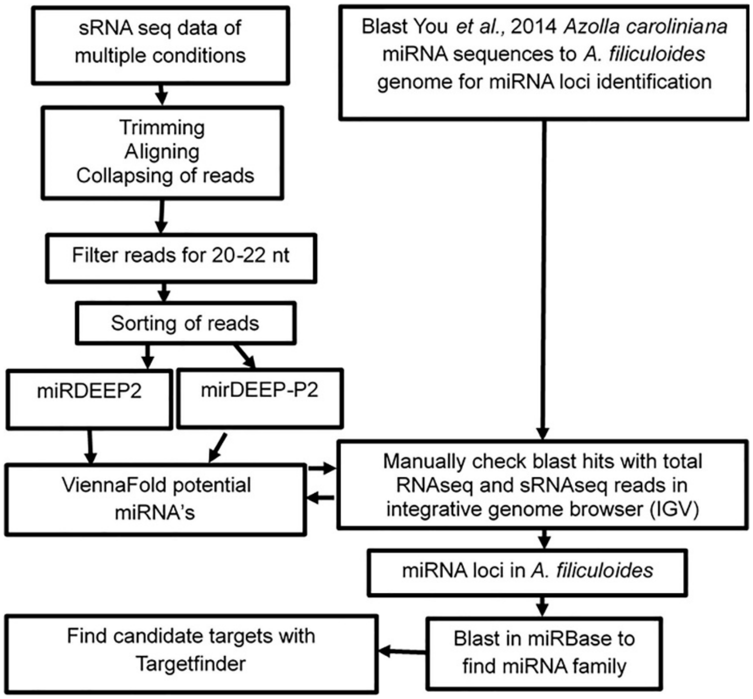

**Supplemental Figure 3**. **Taxonomy of the *Azolla* strains in this study.**

**(A)** Sequence variability of ITS1 intergenic regions of rRNA within the genome of each *Azolla* species (Dijkhuizen *et al.*, 2018). ITS1 sequences were extracted by alignment to the *A. filiculoides* genome (Li *et al*., 2018). Alignments were visualized using WebLogo: at each position of the ITS, the height of the stack indicates the sequence conservation when compared to the *A. filiculoides* reference, while the height of symbols within the stack indicates the relative frequency of each base within the genome of the species analysed.

**(B)**, The typical pointed, two-celled, papillae from the dorsal side of leaves of the Anzali compared to the Galgenwaard strain. Z-stack images were taken of leaves from mature ferns using the Zeiss Axiozoom V.16 binocular with CL9000 LED lights, then reconstituted with Helicon focus software (B, radius 25 smoothing 25).

A

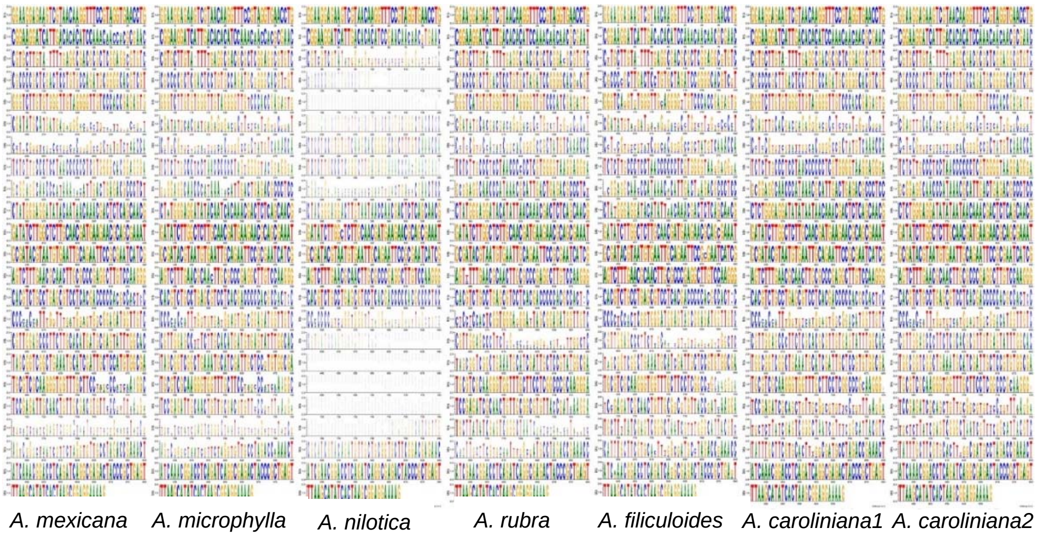


B

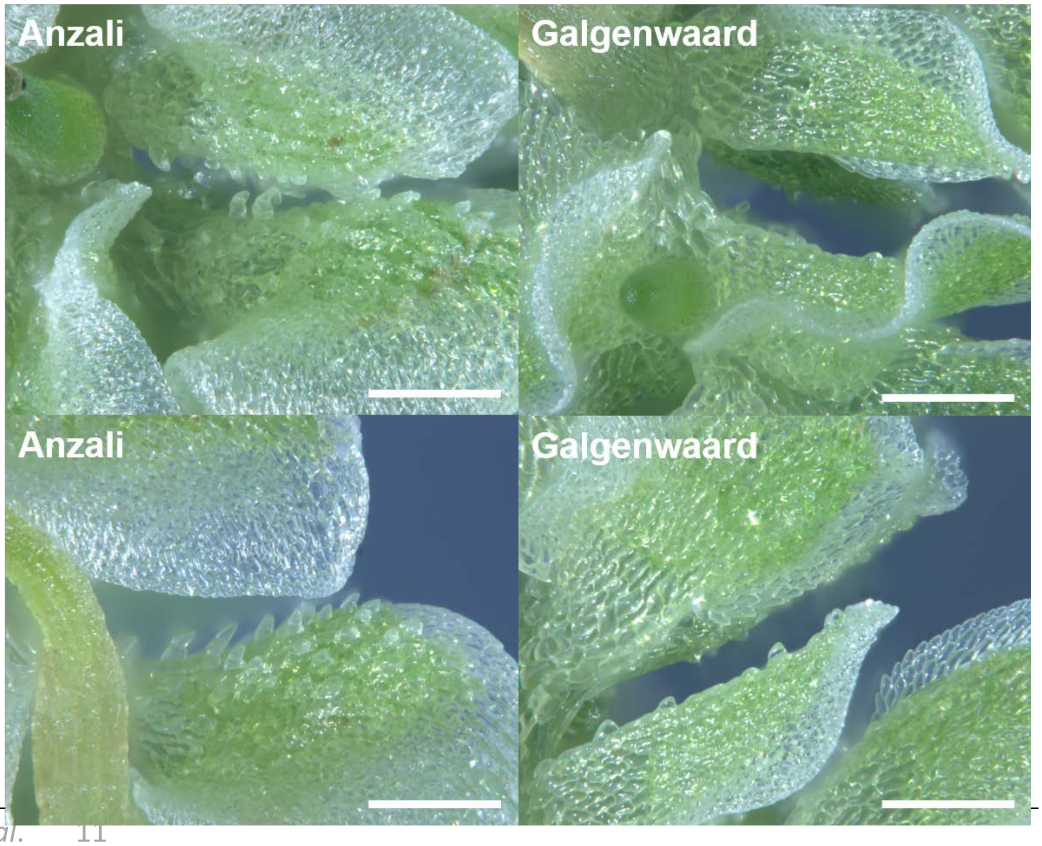

**Supplemental Figure 4.** ***Azolla* MIKC$^C$ phylogenetic analysis and response to far-red light (FR).** The *Azolla* MIKC$^C$ gene model encoded by Azfi_s0028.g024032 was annotated manually: the TF was not expressed in sporophytes yet automated annotation relied on sporophyte RNAsequencing. Sequences extracted from the genome browsers of each species were aligned with MAFFT einsi (Katoh *et al*., 2019), then trimmed with trimAl (Capella-Gutiérrez *et al*., 2009). Phylogenetic inferences were computed with IQTREE (Nguyen *et al*., 2015) and its internal model fitter; 2000 bootstrap determined by SH-aLRT (Kalyaanamoorthy et al., 2017). The tree was rooted using the sequence of Chara globulosa MIKC$^C$ (cgMADS1). Nodes with bootstrap support equalor greater than 80% SH-aLRT are indicated. The lower branches of the tree were typically poorly supported by bootstrapping. Branches are color coded as per their plant lineage. Fold-change inresponse to FR, AND Base Mean were calculated by DESeq2 (Love *et al*., 2014); yellow starsmark significant changes of Padj <0.1.

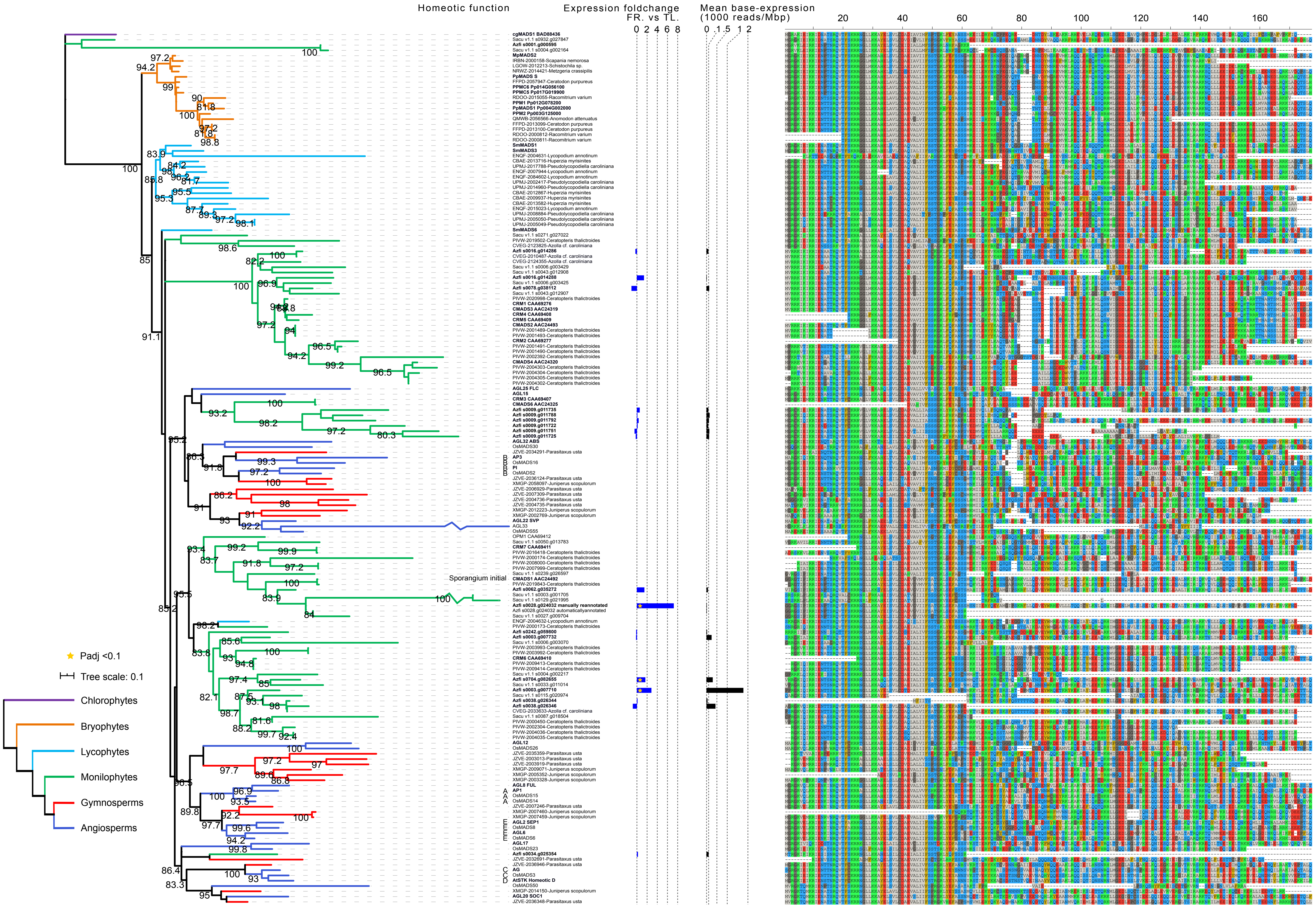

**Supplemental Figure 5. sRNA sequencing and mapping statistics.** sRNA was extracted from sporophytes grown without nitrogen in the medium under TL light with far-red LED (F1-3), and without far red LED (T1-3); sporophytes grown with nitrogen under TL light with far-red LED (FN1-3) and sporophytes depleted of *N.azollae* (C1-3).

**(A)** Yield of quality sRNA reads;

**(B)** Proportion of reads mapping to individual genomes of *A. filiculoides* (Azfi), its chloroplast, *N. azollae* and an associated bacterium from the *Shinella* genus (Shinella) as well as *E.coli* control;

**(C)** sRNA mapping on the concatenated genomes of *A. filiculoides* in (**B**), comparing uniquely mapping versus multi-mappers.

**(D)** Dispersion when comparing 20-22 nt fern nucleus sRNA in sporophytes on TL with and without far-red LED.

**(E)** DESeq2 on the 20-22 nt from (d), red dots represent sRNA with Padj < 0.1.

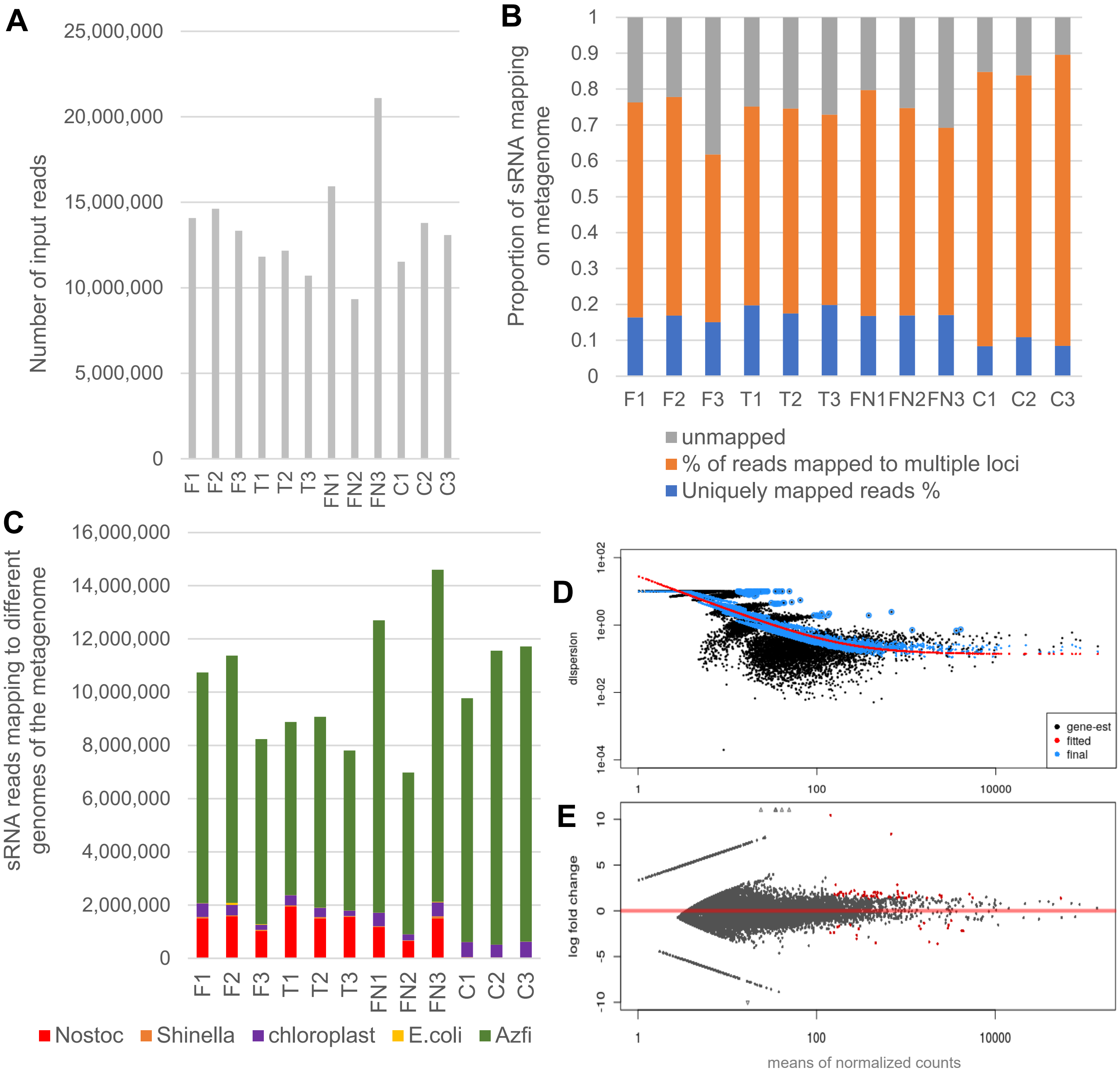

**Supplemental Figure 6. MiR172 target sites in AP2/TOE1 transcription factors (eaAP2 lineage) comparing *A. filiculoides* and seed plants.**

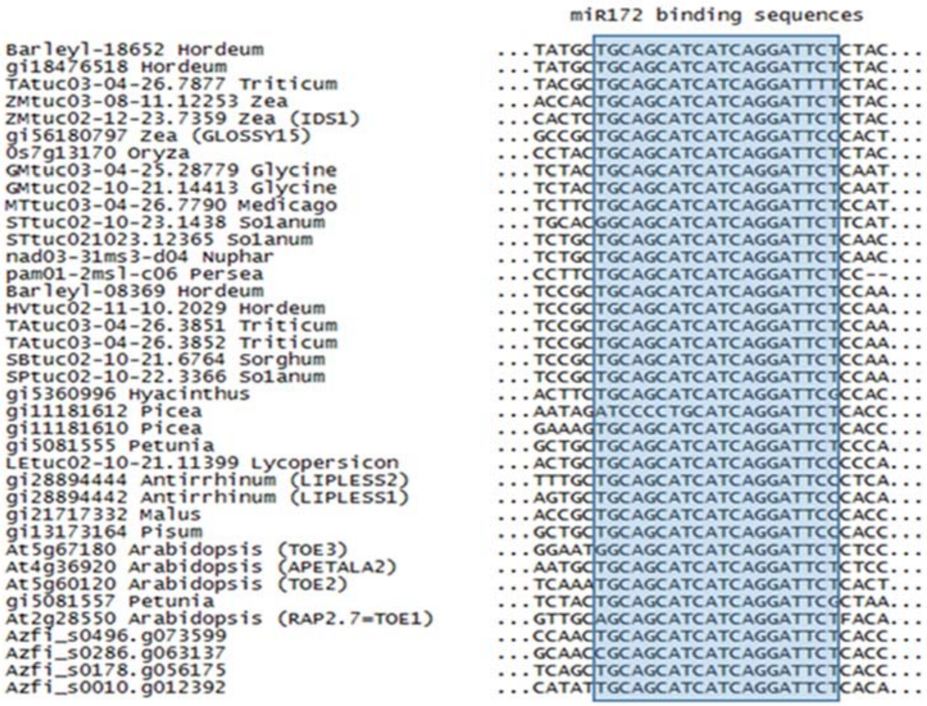

**Supplemental Figure 7. The miRNA156α locus in *A. filiculoides*.** Alignments are visualized using the Integrative Genomics Viewer (Thorvaldsdóttir et al., 2013).
**(A)** Overview of the locus with the 143 b loop.
**(B)** Detail of alignments at the miRNA sequence.
**(C)** Detail of alignments at the miRNA* sequence. mRNA enriched RNA-seq reads were pooled from sporophytes grown under various conditions, harvested at different time points in the diel cycle (Brouwer et al., 2017) and are shown under "RNAseq all conditions". Small-(s)RNA reads were from sporophytes grown under different conditions: without *N. azollae* in far-red light supplemented tube light (sRNAseq –*N. azollae*), with *N. azollae* in far-red light supplemented tube light (sRNAseq +FR) and with *N. azollae* in tube light (sRNAseq –FR). S, read summary; i, individual reads. Azfi_miRNA156α
(Purple box), miRNA* (Pink box). Amino acid sequences in all three reading frames are shown along with boxed stop (red) and start codons (green). Predicted exon (thick blue bar) from automated annotation at Azfi_s0072.g037014 (Li et al., 2018).
**(D)** Azfi_miRNA156α hairpin folded using Vienna RNAfold (Gruber et al., 2008, predicts -79.74 kcal mol$^{-1}$ released at 22 °C), miRNA (purple) and miRNA* (pink).

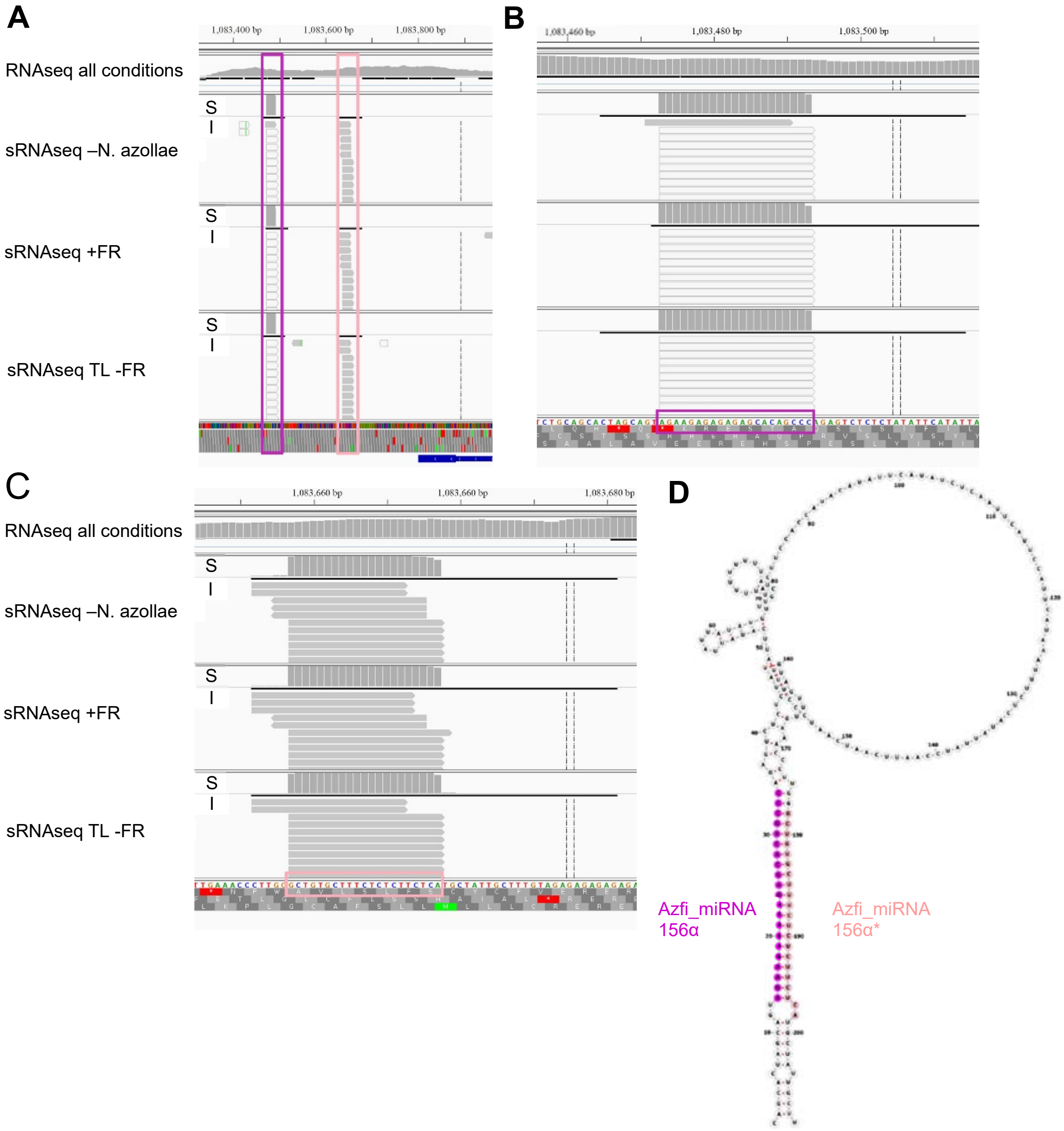

**Supplemental Figure 8. The *AzfiGAMYB* locus Azfi_s0004.g008455 of *A.filiculoides* targeted by miRNA319.**

**(A)** Azfi_miRNA319 hairpin folded using Vienna RNAfold (Gruber et al., 2008, predicts -145,93 kcal $mol^{-1}$ at 37 $^{o}$C); miRNA (purple) and miRNA* (pink).

**(B)** Alignment of reads at the locus.

**(C)** Detail of miRNA319 alignment at the miRNA319 target sequence (red box). S, read summary; I, individual reads. RNAseq all conditions, all reads from Brouwer *et al.*, 2017. Dual RNAseq +FR, reads from sporophytes with FR; Dual RNAseq -FR, reads from sporophytes without FR. Predicted exon (thick blue bar) from automated annotation (Li *et al.*, 2018).

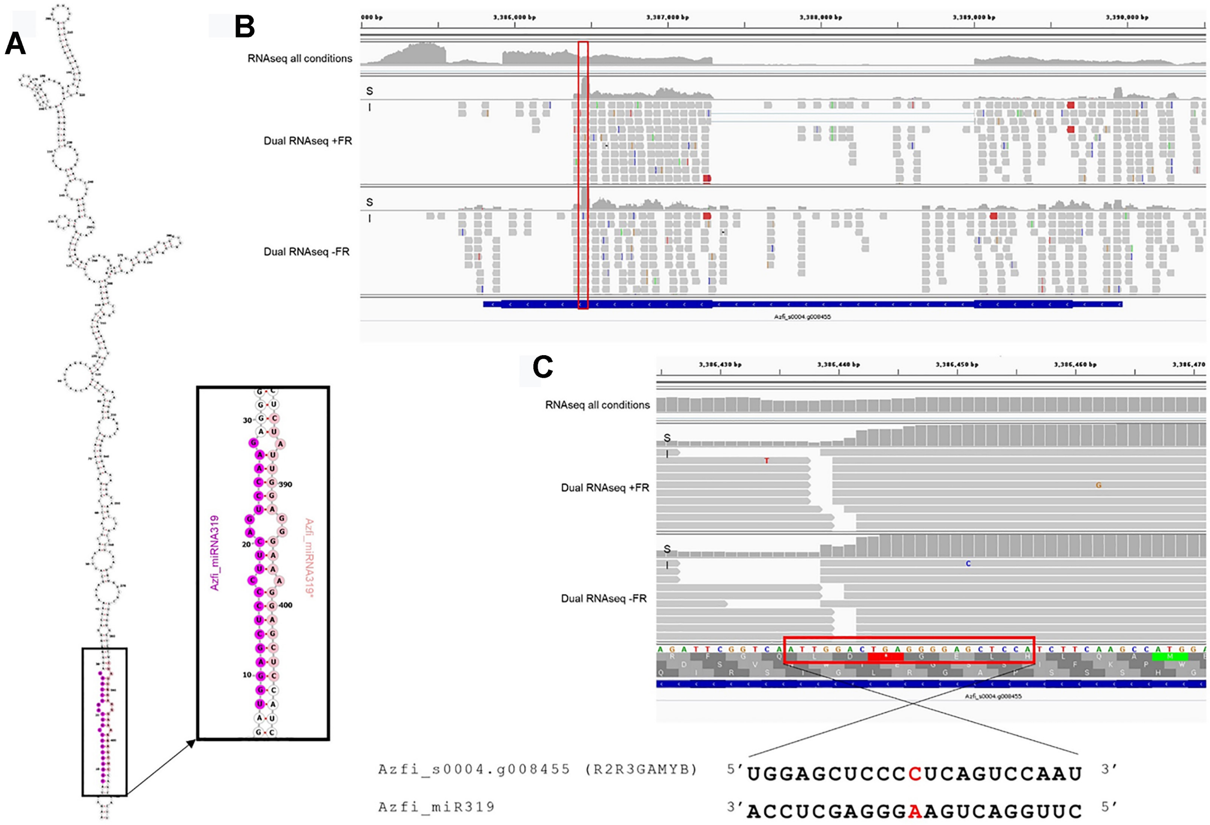

**References Supplemental Data**


**Bowman JL, Kohchi T, Yamato KT, Jenkins J, Shu S, Ishizaki K, Yamaoka S, Nishihama R, Nakamura Y, Berger F, *et al.* 217**. Insights into land plant evolution garnered from the *Marchantia polymorpha* genome. *Cell* **171**:287-34.

**Brouwer P, Brutigam A, Klahoglu C, Tazelaar AOEAOE, Kurz S, Nierop KGJKGJ, van der Werf A, Weber APMAPM, Schluepmann H**. **214**. *Azolla* domestication towards a biobased economy? *New Phytologist* **22**: 169–182.

**Capella-Gutirrez S, Silla-Martnez JM, Gabaldn T. 29.** trimAl: a tool for automated alignment trimming in large-scale phylogenetic analyses. *Bioinformatics* **25**: 1972-3.

**Crooks GE, Hon G, Chandonia JM, Brenner SE**. **24**. WebLogo: A sequence logo generator. *Genome Research* **14**: 1188-119.

**Dai X, Zhuang Z, Zhao PX**. **218**. psRNATarget: a plant small RNA target analysis server (217 release). *Nucleic Acids Research* **46**: W49–W54.

**Dijkhuizen LW, Brouwer P, Bolhuis H, Reichart G-J, Koppers N, Huettel B, Bolger AM, Li F-W, Cheng S, Liu X, *et al.* 218**. Is there foul play in the leaf pocket? The metagenome of floating fern Azolla reveals endophytes that do not fix $N_2$ but may denitrify. *New Phytologist* **217**: 453-466.

**Eren AM, Esen C, Quince C, Vineis JH, Morrison HG, Sogin ML, Delmont TO. 215.** Anvi'o: an advanced analysis and visualization platform for 'omics data. *PeerJ*. **3**:e1319.

**Fahlgren N and Carrington JC. 21.** miRNA target prediction in plants. *Methods in Molecular Biology* **592**: 51– 57.

**Gruber AR, Lorenz R, Bernhart SH, Neubock R, Hofacker IL**. **28**. The Vienna RNA Websuite. *Nucleic Acids Research* **36**: W7–W74.

**Katoh K, Rozewicki J, Yamada KD**. **219.** MAFFT online service: multiple sequence alignment, interactive sequence choice and visualization. *Briefings in bioinformatics* **2**: 116-6.

**Kalyaanamoorthy S, Minh BQ, Wong TK, von Haeseler A, Jermiin LS. 217.** ModelFinder: fast model selection for accurate phylogenetic estimates. *Nature methods*. **14**:587-9.

**Kozomara A, Griffiths-Jones S. 214.** miRBase: annotating high confidence microRNAs using deep sequencing data. *Nucleic acids research* **42**: D68-73.

**Kuang Z, Wang Y, Li L, Yang X**. **219**. miRDeep-P2: accurate and fast analysis of the microRNA transcriptome in plants (I Birol, Ed.). *Bioinformatics* **35**: 2521–2522.

**Kumar S, Stecher G, Tamura K**. **216**. MEGA7: Molecular Evolutionary Genetics Analysis Version 7. for Bigger Datasets. *Molecular biology and evolution* **33**:187-74.

**Kurtz S, Phillippy A, Delcher AL, Smoot M, Shumway M, Antonescu C, Salzberg SL. 24.** Versatile and open software for comparing large genomes. *Genome biology* **5**:R12.

**Leebens-Mack JH, Barker MS, Carpenter EJ, Deyholos MK, Gitzendanner MA, Graham SW, Grosse I, Li Z, Melkonian M, Mirarab S, *et al.* 219**. One thousand plant transcriptomes and the phylogenomics of green plants. *Nature* **574**: 679.

**Letunic I, Bork P**. **219**. Interactive Tree Of Life (iTOL) v4: recent updates and new developments. *Nucleic acids research* **47**: W256-259.

**Li F-W, Brouwer P, Carretero-Paulet L, Cheng S, de Vries J, Delaux P-M, Eily A, Koppers N, Kuo L-Y, Li Z, *et al.* 218**. Fern genomes elucidate land plant evolution and cyanobacterial symbioses. *Nature Plants* **4**: 46-472.

**Li H, Durbin R**. **2009**. Fast and accurate short read alignment with Burrows-Wheeler transform. *Bioinformatics* **25**: 1754–1760.
**Madeira PT, Hill MP, Dray FA, Coetzee JA, Paterson ID, Tipping PW**. **2016**. Molecular identification of Azolla invasions in Africa: The Azolla specialist, Stenopelmus rufinasus proves to be an excellent taxonomist. *South African Journal of Botany* **105**: 299-305.
**von Meijenfeldt FB, Arkhipova K, Cambuy DD, Coutinho FH, Dutilh BE. 2019.** Robust taxonomic classification of uncharted microbial sequences and bins with CAT and BAT. *Genome biology* **20**:217.
**Nguyen LT, Schmidt HA, Von Haeseler A, Minh BQ. 2015.** IQ-TREE: a fast and effective stochastic algorithm for estimating maximum-likelihood phylogenies. *Molecular biology and evolution* **32**: 268-74.
**Nurk S, Meleshko D, Korobeynikov A, Pevzner PA. 2017**. metaSPAdes: a new versatile metagenomic assembler. *Genome research* **27**:824-34.
**Olm MR, Brown CT, Brooks B, Banfield JF. 2017.** dRep: a tool for fast and accurate genomic comparisons that enables improved genome recovery from metagenomes through de-replication. *The ISME Journal* **11**: 2864–2868.
**Slater GS, Birney E. 2005.** Automated generation of heuristics for biological sequence comparison. *BMC bioinformatics **6**:31.*
**Thompson JD, Higgins DG, Gibson TJ**. **1994**. CLUSTAL W: Improving the sensitivity of progressive multiple sequence alignment through sequence weighting, position-specific gap penalties and weight matrix choice. *Nucleic Acids Research* **22**:4673-80.
**Thorvaldsdóttir H, Robinson JT, Mesirov JP. 2013.** Integrative Genomics Viewer (IGV): high-performance genomics data visualization and exploration. *Briefings in bioinformatics*. **14**: 178-92.
**Trifinopoulos J, Nguyen LT, von Haeseler A, Minh BQ**. **2016**. W-IQ-TREE: a fast online phylogenetic tool for maximum likelihood analysis. *Nucleic acids research* **44**: W232-235.
